# Cortex Folding by Combined Progenitor Expansion and Adhesion-Controlled Neuronal Migration

**DOI:** 10.1101/2024.05.08.593109

**Authors:** Seung Hee Chun, D. Santiago Diaz Almeida, Mihail Ivilinov Todorov, Tobias Straub, Tobias Ruff, Wei Shao, Jianjun Yang, Gönül Seyit-Bremer, Yi-Ru Shen, Ali Ertürk, Daniel del Toro, Songhai Shi, Rüdiger Klein

## Abstract

Folding of the mammalian cerebral cortex into sulcal fissures and gyral peaks is the result of complex processes that are incompletely understood. Previously we showed that genetic deletion of Flrt1/3 adhesion molecules causes folding of the smooth mouse cortex into sulci resulting from increased lateral dispersion and faster neuron migration, without progenitor expansion. Here, we find that combining the *Flrt1/3* double knockout with an additional genetic deletion that causes progenitor expansion, greatly enhances cortex folding. Expansion of intermediate progenitors by deletion of Cep83 results in enhanced formation of sulci. Expansion of apical progenitors by deletion of Fgf10 results in enhanced formation of gyri. Single cell transcriptomics and simulations suggest that changes in adhesive properties of cortical neurons, their proportions and densities in the cortical plate, combined with lateral dispersion during their radial migration are important folding parameters. These results identify key developmental mechanisms that cooperate to promote cortical gyrification.

**HIGHLIGHTS:**

- Cortex folding is enhanced by combining progenitor expansion and divergent migration.
- Concomitant expansion of intermediate progenitors results in the formation of sulci
- Concomitant expansion of apical progenitors results in the formation of gyri
- Progenitors differentially affect cortical neurons with distinct adhesive properties

## INTRODUCTION

One of the most remarkable aspects of brain development is the folding of the cerebral cortex, which is believed to enhance the cognitive capacities of large mammals. Brain size has changed dramatically during mammalian evolution^1^ and this is mostly due to an increase in size of the cerebral cortex. The cerebral cortex is highly folded into peaks (gyri) and fissures (sulci) in gyrencephalic species, or has a smooth surface in lissencephalic, typically smaller mammals such as the mouse.^2^

Previous studies have identified mechanisms that drive cortex folding, including cell proliferation, migration and biomechanical properties of the tissue.^3^ Most studies have focused on amplification of cell proliferation due to the correlation between larger brains and cortical folding.^4,5^ Progenitors are distributed in two different cortical germinal layers: the ventricular zone (VZ), which includes the neuroepithelial cells (NECs) and apical radial glia (aRG), and the subventricular zone (SVZ) containing intermediate progenitors (IPs) and basal radial glia (bRG).^3^ A massive increase in the surface area of both layers is an evolutionary hallmark of gyrencephalic species. This includes the thickening of the SVZ, which is then subdivided into an inner (ISVZ) and an outer (OSVZ) subventricular zone.^6,7,8^ The SVZ in gyrencephalic species like the ferret is not homogeneous as in the mouse, and displays microdomains with high and low proliferation of progenitors, a pattern that favors the formation of future gyri and sulci, respectively.^9^ Supporting this model, local amplification of progenitors in the murine cortex induces the formation of gyrus-like structures.^10–13^ Interestingly, studies inducing uniform progenitor expansion in the mouse cortex have observed thickening of this structure without folding.^14,15^ These results suggest that progenitor expansion in discrete cortical domains is a key event to induce gyration of the cerebral cortex.

Generating genetic mouse models in which cortex folding is induced by progenitor expansion has remained a challenge. Elevated Sonic hedgehog (Shh) signaling by expressing a constitutively active receptor^16^, or increased Wnt signaling through the loss of transcription factors Lmx1a/1b^17^, lead to IP expansion and cortex folding in mice. However, these modifications affect the survival of mutant embryos at later developmental and postnatal stages. One exception is the genetic ablation of the centrosomal protein 83 (Cep83). Its deletion removes the apical membrane anchoring of the centrosome in aRGs and induces an expansion of IPs. This progenitor amplification mainly occurs in the anteromedial regions of the cortex, which correlates with cortex folding.^18^

Fibroblast growth factor (FGF) signaling is one of the most relevant pathways that regulate cortical proliferation, in addition to Wnt, Shh and Notch.^19^ Indeed, the expression of a constitutively active Fgf3 mutant allele (K664E) in mice, leads to an increase in cortical thickness, but not cortex folding.^20^ Ventricular injection of FGF2 leads to increased tangential growth of the VZ and proliferation of IP that correlates with cortex folding.^21^ Similarly, local induction of FGF8 in the cortex increases the number of IPs and the formation of additional gyri in the ferret.^22^ Conversely, suppressing all FGFR activity in the ferret cortex impairs the formation of cortical folds.^23^ Ablation of Fgf2 in mice leads to reduced expansion and thickening of the cortex^24^, but this phenotype is opposite to that in Fgf10 knockout mice.^25^ FGF10 is expressed mainly by rostral cortical NECs during embryonic day 9 (E9) to E11, where it supports their differentiation into aRGs.^26^ Genetic deletion of Fgf10 delays their differentiation, favoring their tangential amplification. This results in an increased number of aRGs and neurons, leading to the expansion and thickening of the cortex in rostral regions.^25^

Besides proliferation, neuronal migration is another key mechanism that drives cortex folding. Indeed, the role of several genes linked to abnormal folding in humans has been associated with neuronal migration in ferret and mouse models.^4,27^ In gyrencephalic species such as the ferret, migrating neurons do not follow strictly parallel pathways as observed in the mouse cortex^28^, but instead, follow divergent trajectories concomitant with the onset of cortical folding.^29^ This is consistent with the observation that neurons in ferret cortices can switch between radial fibers and enhance their lateral dispersion, compared to those in the mouse.^30^ We provided a molecular mechanism for the role of neuronal migration in cortex folding, demonstrating that genetic knockout of Flrt1/3 adhesion molecules leads to faster and divergent migration, as well as sulcus formation, in the mouse cortex without changes in cell proliferation.^31^ This phenotype arises from changes in the intercellular adhesion among migrating neurons, where approximately 50% are FLRT-mutant neurons (previously destined to become FLRT+ in wild-type animals). These FLRT-mutant neurons exhibit reduced intercellular adhesion, migrate faster, and segregate from their neighboring FLRT-negative neurons. A similar scenario has been observed for the Eph/ephrin family of guidance receptors/ligands. Overexpression of ephrinB1, which can induce homophilic cell adhesion^32^,reduces the horizontal dispersion of multipolar neurons.^33^ Likewise, EphA/ephrinA gain-of-function experiments show reduced lateral dispersion of multipolar neurons.^34^ Similarly, ephrinA2/A3/A5 triple knockout (tKO) mice, like the *Flrt1/3* dKO phenotype, display neuronal segregation along the tangential axis, leading to a wavy cortical plate with alternating thicker and thinner areas.^34^

A widely accepted hypothesis suggests that cortex folding is driven by a combination of uneven progenitor expansion and divergent radial migration.^35^ Yet, we still do not fully understand the logic behind how coordinated and simultaneous events like proliferation and divergent migration lead to the formation of sulci and gyri in the cortex. Here, we addressed this question by taking advantage of the *Flrt1/3* dKO mouse line, which promotes divergent migration, and combined it with other genetic lines that induce progenitor expansion. We chose two different lines that expand either intermediate or apical progenitors through genetic ablation of Cep83 and Fgf10, respectively. This allowed us to dissect the specific contributions of distinct progenitor pools to cortex folding when coupled with increased divergent migration. We found that expansion of intermediate progenitors favors sulcus formation, whereas amplification of apical progenitors promotes gyrus formation in the *Flrt1/3* dKO model. Single cell transcriptomics and simulations suggest that changes in adhesive properties of cortical neurons, cell sorting by differential adhesion, increased cell densities in the cortical plate, combined with lateral dispersion during their radial migration are important folding parameters. Our results identify key developmental mechanisms that collaboratively drive cortex folding.

## RESULTS

### Loss of Cep83 and Flrt1/Flrt3 enhances cortical folding and the presence of sulci

To investigate the contributions of progenitor cell expansion and divergent cell migration to cortex folding, we combined the deletion of Cep83 with the double deletion of Flrt1 and Flrt3. In previous work^31^ we had used the pan-nervous system Nestin-Cre driver to conditionally delete Flrt3 and, together with the Flrt1 null allele, to generate *Flrt1/3* double knockout (dKO) mice. The conditional deletion of Cep83 was previously done with the Emx1-Cre driver^18^ which is expressed predominantly in the developing cortex and hippocampus starting at E10.5 with an earlier developmental time course than Nestin-Cre (Figure S1 A,B). We decided to use the Emx1-Cre driver to generate *Cep83;Flrt1/3* triple KO mice (from now on referred to as *Cep83* tKO mice). We validated that the Emx1-Cre driver was effective in deleting Cep83 and FLRT3 protein expression (Figure S1 C,D) and that the phenotype of *Emx1-Cre*;*Flrt1/3* dKO mice (from now on referred to as Emx1-*Flrt1/3* dKO mice) was overall very similar to *Nestin-Cre*;*Flrt1/3* dKO mice. At E17.5, the size of the *Flrt1/3* dKO cortex was not enlarged compared to *Flrt1*^−/−^*Flrt3*^lx/+^ mice or heterozygous controls, consistent with the previous observation that progenitor populations were not expanded (Figure 1 A and FigureS1 E and data not shown). Approx. one third of the animals had folds, often only a single fold, and typically in the lateral cortex (Figure 1B). Occasionally, we also observed folds in the lateral region of the cortex in control littermates, deficient for Flrt1, in line with previous findings^31^ (Figure S1 F). We also analyzed cortex size and folding of E17.5 *Emx1-Cre*;*Cep83* single cKO embryos, and found that the cortex was expanded as expected (Figure 1 C and Figure S1 G) and folds were observed in the medial (cingulate) cortex as previously described for P21 mutants^18^ (Figure S1 H). We then generated *Cep83* tKO mice and found that cortex sizes of E17.5 embryos were enlarged (Figure 1 D,E) and, more interestingly, that the numbers of cortical folds were more numerous than in the *Cep83* single and *Flrt1/3* dKO mice combined. Typically, cortical folds could be seen in freshly dissected brains and after tissue clearing and light sheet imaging (Figure 1,G; movie 1,2). Folds were often seen as individual sulci in lateral and/or cingulate cortex, either unilateral or bilateral (Figure 1 H-J). Cortical layers were preserved as shown by immunostainings for upper layer (Satb2) and deeper layer neurons (Ctip2) (Figure 1 H-J). The apical surface of the ventricular zone remained unfolded, indicating *bona fide* foldings^6^ (Figure 1 H-J). Importantly, the penetrance of the phenotype was greatly increased from approx. one third in *Cep83* single and *Flrt1/3* dKO mice to over 80% penetrance in *Cep83* tKO mice (Figure 1 K,L). Remarkably, mice homozygous for Cep83 and Flrt1 deletion, but heterozygous for Flrt3 (*Emx1-Cre*;*Cep83*^fl/fl^;*Flrt1*^−/−^;*Flrt3*^fl/+^ double KO (*Cep83* dKO)) also showed increased penetrance of cortical folding (Figure 1 K,L). This was consistent with previous observations in Nestin-Cre driven *Flrt1/3* mutant mice.^31^ We calculated the gyrification index (GI) (a measure of the real length of the cortical surface over a hypothetical minimal length of the cortical surface) and found that in rostral and medial regions of the *Cep83* tKO cortex, the GI was significantly higher than in single *Cep83* cKO or *Emx1-Flrt1/3* dKO mice (Figure 1 M,N). These results indicate that deletion of Cep83 together with deletion of Flrt1/3 leads to a strong cortex folding phenotype with the presence of bilateral sulci.

**Figure 1.**
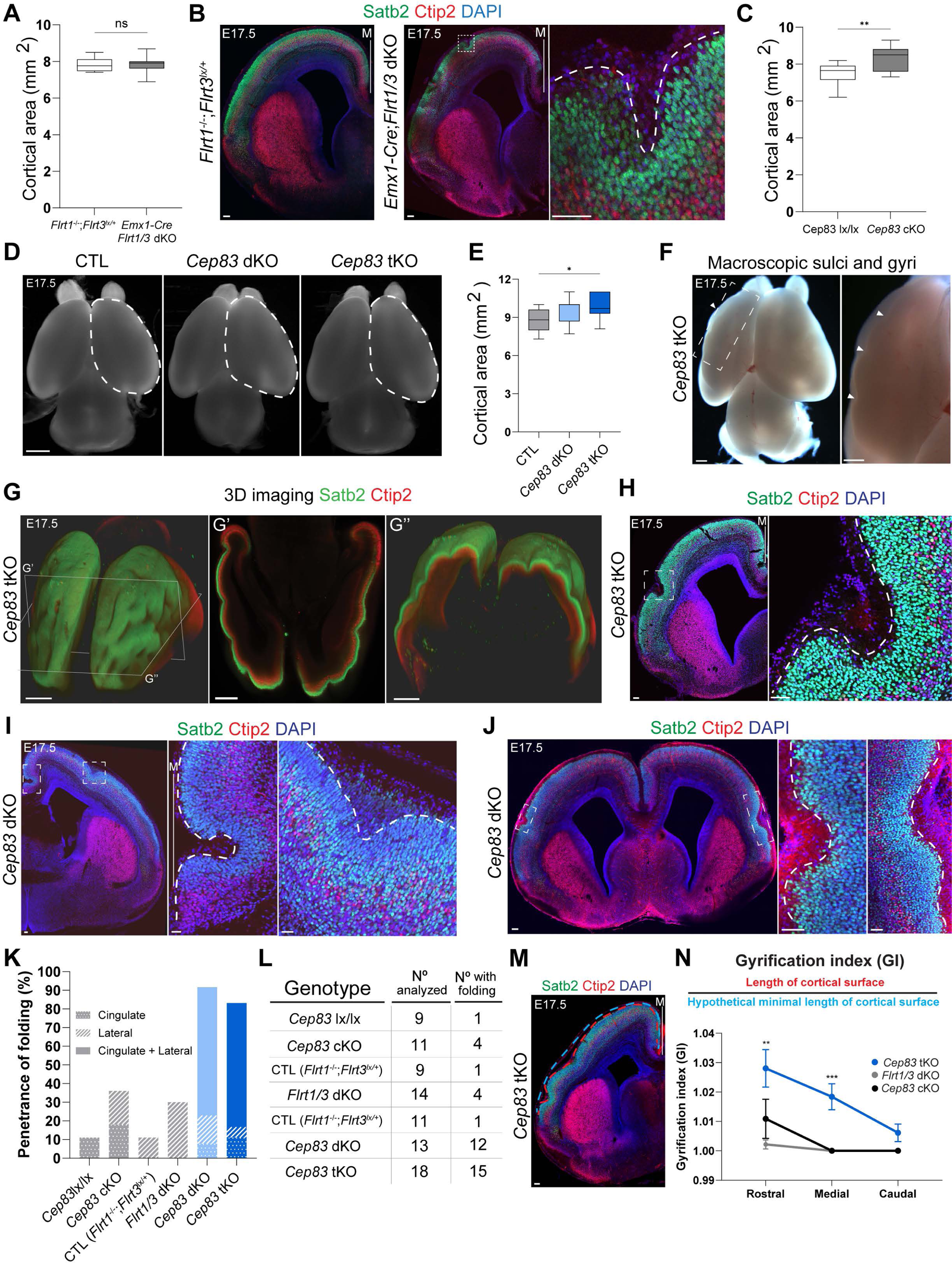
Loss of Cep83 and Flrt1/Flrt3 enhances cortical folding and the presence of sulci. (A) Quantification of cortical areas. Data are represented as a box plot, with median (center line), interquartile range (box) and minimum and maximum values (whiskers); *Flrt1* ^−/−^*Flrt3* ^lx/+^, n = 9 brains; *Emx1-Cre*;*Flrt1/3* dKO, n = 13 brains. p = 0.916, t-test with Welch correction. (B) E17.5 *Flrt1* ^−/−^*Flrt3* ^lx/+^ and *Emx1-Cre*;*Flrt1/3* dKO brain sections labeled with Satb2 (green), Ctip2 (red), and DAPI (blue). Area in dashed rectangle is shown with higher magnification on the right. The midline is indicated with a vertical line and the letter ‘M’. (C) Quantification of cortical areas. *Cep83* lx/lx, n = 14 brains; *Cep83* cKO, n = 16 brains. p = 0.002, **p < 0.01, t-test with Welch correction. (D) Representative whole-mount images of E17.5 CTL (*Flrt1*^−/−^;*Flrt3*^lx/+^), *Cep83* dKO, and *Cep83* tKO brains. Dashed areas were measured to obtain quantification in (E). (E) Quantification of the cortical areas. CTL (*Flrt1*^−/−^;*Flrt3*^lx/+^), n = 11 brains; *Cep83* dKO, n = 13 brains; *Cep83* tKO n = 19 brains. p = 0.021, *p < 0.05, one-way ANOVA, Tukey’s post hoc analysis. (F) Macroscopic sulci and gyri in an E17.5 *Cep83* tKO embryo. The area in the dashed rectangle is shown with higher magnification on the right, sulci and gyri are indicated by arrowheads. (G-G’’) 3D imaging of the E17.5 *Cep83* tKO embryo shown in (F) labeled with Satb2(green) and Ctip2(red). 3D whole brain, XY plane and YZ plane single images are shown (whole dataset in Movie 1). (H) E17.5 *Cep83* tKO brain section labeled with Satb2 (green), Ctip2 (red), and DAPI (blue). A sulcus in a dashed rectangle is shown with higher magnification on the right. The midline is indicated with a vertical line and the letter ‘M’. (I) E17.5 *Cep83* dKO brain section labeled with Satb2 (green), Ctip2 (red), and DAPI (blue). Two sulci in cingulate and lateral cortex in dashed rectangles are shown with higher magnification on the right. The midline is indicated with a vertical line and the letter ‘M’. (J) E17.5 *Cep83* dKO brain section labeled with Satb2 (green), Ctip2 (red), and DAPI (blue). Bilateral sulci in dashed rectangles are shown with higher magnification on the right. (K) Folding penetrance in E17.5 *Cep83* lx/lx (littermate CTL of *Cep83* cKO), *Cep83* cKO, CTL (*Flrt1*^−/−^;*Flrt3*^lx/+^), littermate controls of *Flrt1/3* dKO and *Cep83* tKO), *Flrt1/3* dKO, *Cep83* dKO, and *Cep83* tKO mice. Locations of sulci are indicated in the graph: Cingulate cortex (dotted pattern), lateral cortex (diagonal pattern), and both (solid fill). (L) Table of cortical folding penetrance at E17.5 for the indicated genotypes. Immunostained brain sections were analyzed for the presence of one or more sulci as shown in Figure 1L. (M) Representative immunostained image of a *Cep83* tKO embryo depicting the quantification of the Gyrification Index (GI). Red line indicates the de facto length of the cortical surface; blue line indicates the hypothetical minimal length of the cortical surface. The midline is indicated with a vertical line and the letter ‘M’. (N) Quantification of GI values. *Cep83* single cKO (black) n = 11 brains, *Flrt1/3* dKO (gray) n = 13 brains, *Cep83* tKO (blue) n = 31 sections from a total of 18 brains, quantified at three positions: rostral, medial, and caudal. Data are shown as mean ± SEM; Rostral : *Cep83* cKO vs. *Flrt1/3* dKO, p = 0.165, *Cep83* cKO vs. *Flrt1/3* dKO p = 0.579, *Flrt1/3* dKO vs. *Cep83* tKO, p = 0.001, Medial : *Cep83* cKO vs. *Flrt1/3* dKO, p > 0.9999, *Cep83* cKO vs. *Flrt1/3* dKO p = 0.009, *Flrt1/3* dKO vs. *Cep83* tKO, p = 0.001, Caudal : *Cep83* cKO vs. *Flrt1/3* dKO, p > 0.9999, *Cep83* cKO vs. *Flrt1/3* dKO p = 0.295, *Flrt1/3* dKO vs. *Cep83* tKO, p = 0.141. **p < 0.01, ***p < 0.001, one-way ANOVA with Tukey’s post hoc analysis. Scale bars represent 100 μm, 100 μm, 100 μm (B), 1 mm (D), 500 μm, 500 μm (F), 300 μm, 400 μm, 400 μm (G-G’’), 100 μm, 50 μm (H), 200 μm, 100 μm. 100 μm (I), 100 μm, 50 μm, 50 μm (J), and 100 μm (M).

### *Cep83* tKO mice have more intermediate progenitors

We next investigated the underlying mechanism of the enhanced folding phenotype of *Cep83* tKO mice. Previous work had indicated an increased density of TBR2+ IPs in the subventricular zone of *Cep83* cKO embryos, whereas the density of mitotic stem cells positive for the marker phosphorylated histone H3 (pH3) at the ventricular zone surface had remained unchanged^18^. We performed quantifications in rostral and medial cortical sections of E13.5 *Cep83* tKO embryos by quantifying the numbers of apical progenitors (Sox2+), IPs (Tbr2+), and mitotic dividing cells (PH3+) (Figure 2 A-C). Moreover, we estimated the number of PH3+ basal cells stained for phospho-vimentin (pVIM), which is known to be expressed in RG cells during cell division.^36^ These basal progenitors are known to promote tangential growth of the cortex with a folded structure in gyrencephalic species^37,38^ (Figure 2 D-F). We used *Flrt1*^−/−^;*Flrt3*^lx/+^ embryos as controls, because they could be generated as littermates and were previously shown to have normal numbers of cortical progenitors^31^. At both rostral and medial location, we observed significantly increased cell numbers of basal IPs (Tbr2+) and double positive Pvim+/PH3+ cells in *Cep83* tKO embryos compared to controls, consistent with previous findings in Cep83 single KO^18^. The total numbers of apical progenitors (Sox2+) and mitotic dividing cells remained unchanged (Figure 2 A-F, S2 A-D). Similar results were obtained with *Cep83* dKO embryos consistent with the observed folding phenotype (Figure 2 A-E). Collectively, these results indicate that deletion of Cep83 causes an expansion of IPs and bRG cells at E13.5 that results in an enlarged cortex at E17.5. Since this effect is already present in *Cep83* cKO mice, it is independent of the presence of functional alleles of Flrt1 and Flrt3. These findings further suggest that the expansion of IPs and bRG cells in *Cep83* tKO embryos exacerbates the cortex folding phenotype seen in the *Flrt1/3* dKO mice.

**Figure 2.**
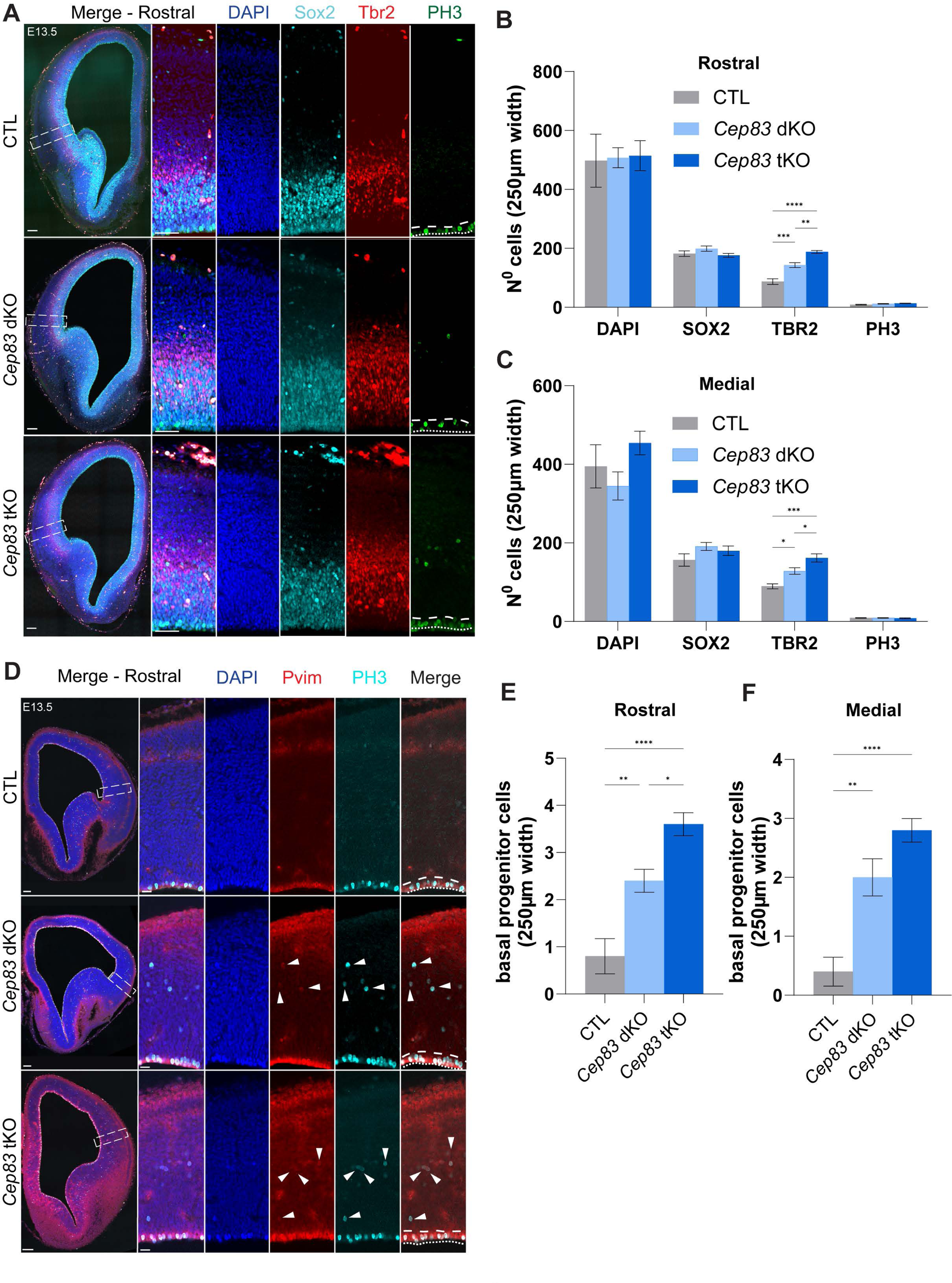
*Cep83* tKO mice have more intermediate progenitors. (A) E13.5 rostral cortices of CTL (*Flrt1*^−/−^;*Flrt3*^lx/+^), *Cep83* dKO. and *Cep83* tKO embryos were stained with DAPI (blue), Sox2 for early neuronal progenitors (cyan), Tbr2 for intermediate progenitors (red), and PH3 for mitotic cells (green). The apical side of the VZ is indicated with a dotted line, the basal side with a dashed line in PH3 stained images. Areas in dashed rectangles in (A, left row) are shown with higher magnification on the right. (B) Quantifications of cell densities in the rostral regions shown in (A). (CTL, *Flrt1*^−/−^;*Flrt3*^lx/+^ n = 5 brains, *Cep83* dKO n = 5 brains, *Cep83* tKO, n = 5 brains). Data are shown as mean ± SEM; CTL vs. *Cep83* dKO p = 0.001, CTL vs. *Cep83* tKO p < 0.001, *Cep83* dKO vs. *Cep83* tKO p = 0.005. **p < 0.01, ***p < 0.001, ****p < 0.0001, one-way ANOVA with Tukey’s post hoc analysis. (C) Quantifications of cell densities in the medial regions shown in Figure S2 A. (CTL, *Flrt1*^−/−^;*Flrt3*^lx/+^ n = 5 brains, *Cep83* dKO n = 5 brains, *Cep83* tKO, n = 5 brains). Data are shown as mean ± SEM; CTL vs. *Cep83* dKO p = 0.019, CTL vs. *Cep83* tKO p = 0.001, *Cep83* dKO vs. *Cep83* tKO p = 0.043. *p < 0.05, ***p < 0.001, one-way ANOVA with Tukey’s post hoc analysis. (D) E13.5 rostral cortices of CTL (*Flrt1*^−/−^;*Flrt3*^lx/+^), *Cep83* dKO. and *Cep83* tKO embryos were stained with DAPI (blue), Pvim for dividing RG cells (red), PH3 for mitotic cells (cyan). The apical side of the VZ is indicated with a dotted line, the basal side with a dashed line in Pvim/PH3 images. The basal progenitors are present as Pvim/PH3 co-immunopositive cells located above apical side of the VZ (marked by arrow heads). (E) Quantifications of cell densities in the rostral regions shown in (D). (CTL, *Flrt1*^−/−^;*Flrt3*^lx/+^ n = 5 brains, *Cep83* dKO n = 5 brains, *Cep83* tKO, n = 5 brains). Data are shown as mean ± SEM; p = 0.034, p = 0.006, *p < 0.05, **p < 0.01, ****p < 0.0001, one-way ANOVA with Tukey’s post hoc analysis. (F) Quantifications of cell densities in the medial regions shown in Figure S2 A. (CTL, *Flrt1*^−/−^;*Flrt3*^lx/+^ n = 5 brains, *Cep83* dKO n = 5 brains, *Cep83* tKO, n = 5 brains). Data are shown as mean ± SEM; p = 0.002, **p < 0.01, ****p < 0.0001, one-way ANOVA with Tukey’s post hoc analysis. Scale bars represent 100 μm, 50 μm (A), and 100 μm, 20 μm (D).

### Loss of Fgf10 and Flrt1/Flrt3 enhances cortical folding and the presence of gyri

We next investigated the interaction of progenitor expansion and cortical cell migration in a separate mouse model, this time involving the deletion of Fgf10. Fgf10 null animals displayed delayed RG differentiation causing enhanced symmetric divisions of neuroepithelial cells early during cortical development eventually resulting in an enlargement of rostral cortical areas at birth^25^. To generate conditional *Fgf10;Flrt1/3* tKO mice (in short *Fgf10* tKO mice), we used the Foxg1-Cre driver, which was previously used in combination with a Fgf10^lx/lx^ allele to enlarge the brain (D. Kawaguchi, Y. Gotoh, unpublished results). Foxg1-Cre expresses uniformly in telencephalic progenitors as early as E9.0-E9.5^39^, one day earlier than Emx1-Cre using the Cre-dependent reporter pCALNL-TdTom (Figure S3 A). We also confirmed that the Foxg1-Cre driver deleted expression of Fgf10 and Flrt3 in the cortex (Figure S3 B-E). We next asked if Foxg1-Cre mice could be used to generate *Flrt1/3* dKO mice. Indeed, we found that *Foxg1-Cre*;*Flrt1/3* dKO mice displayed a cortex folding phenotype that was qualitatively similar to *Nestin-Cre*;*Flrt1/3* dKO mice (Figure S3 F), consisting mostly of a single sulcus, albeit with very low penetrance (see below). Next, we investigated the effects of Fgf10 deletion on cortex size. As expected, we found that the absence of Fgf10 increased rostral cortical thickness compared to controls lacking Foxg1-Cre (Figure 3 A,B). In few embryos this phenotype correlated with a gyrus-like protrusion (Figure 3C). This effect was not observed in *Foxg1-Cre*;*Flrt1/3* dKO mice with an intact Fgf10 gene (Figure 3 D,E). Next, we generated *Fgf10* tKO mice and, by measuring the entire size of the cortex, found that it was expanded (Figure 3 F,G). This was unexpected, since using the same measurements neither *Fgf10* single cKO nor *Flrt1/3* dKO mice had an expanded cortex (Figure 3 H,I, S3G,H). Interestingly, *Fgf10* tKO mice showed enhanced and bilateral cortex folding, but unlike *Cep83* tKO mice, folds more often resembled gyri (Figure 3 J-L). Overall, cortical layers were preserved as indicated by the markers Satb2 and Ctip2, and the apical surface of the VZ remained unaltered, indicating a *bona fide* cortical folding^6^ phenotype (Figure 3 K,L). Penetrance was very much enhanced from approx. 10-15% in *Fgf10* single cKO and *Flrt1/3* dKO mice to over 90% penetrance in *Fgf10* tKO mice (Figure 3 M,N). The folding phenotype was also highly penetrant in mice homozygous for Fgf10 and Flrt1 deletion, and heterozygous for Flrt3 (*Foxg1-Cre*;*Fgf10*^fl/fl^;*Flrt1*^−/−^;*Flrt3*^fl/+^ double KO [in short *Fgf10* dKO]) (Figure 3M,N), consistent with the *Cep83* dKO situation). The GI was significantly higher in Flrt1/3 tKO than *Fgf10* single cKO mice (Figure 3 O,P). These results indicate that deletion of Fgf10, combined with Flrt1/3 ablation, leads to a stronger cortex folding phenotype with the presence of bilateral gyri.

**Figure 3.**
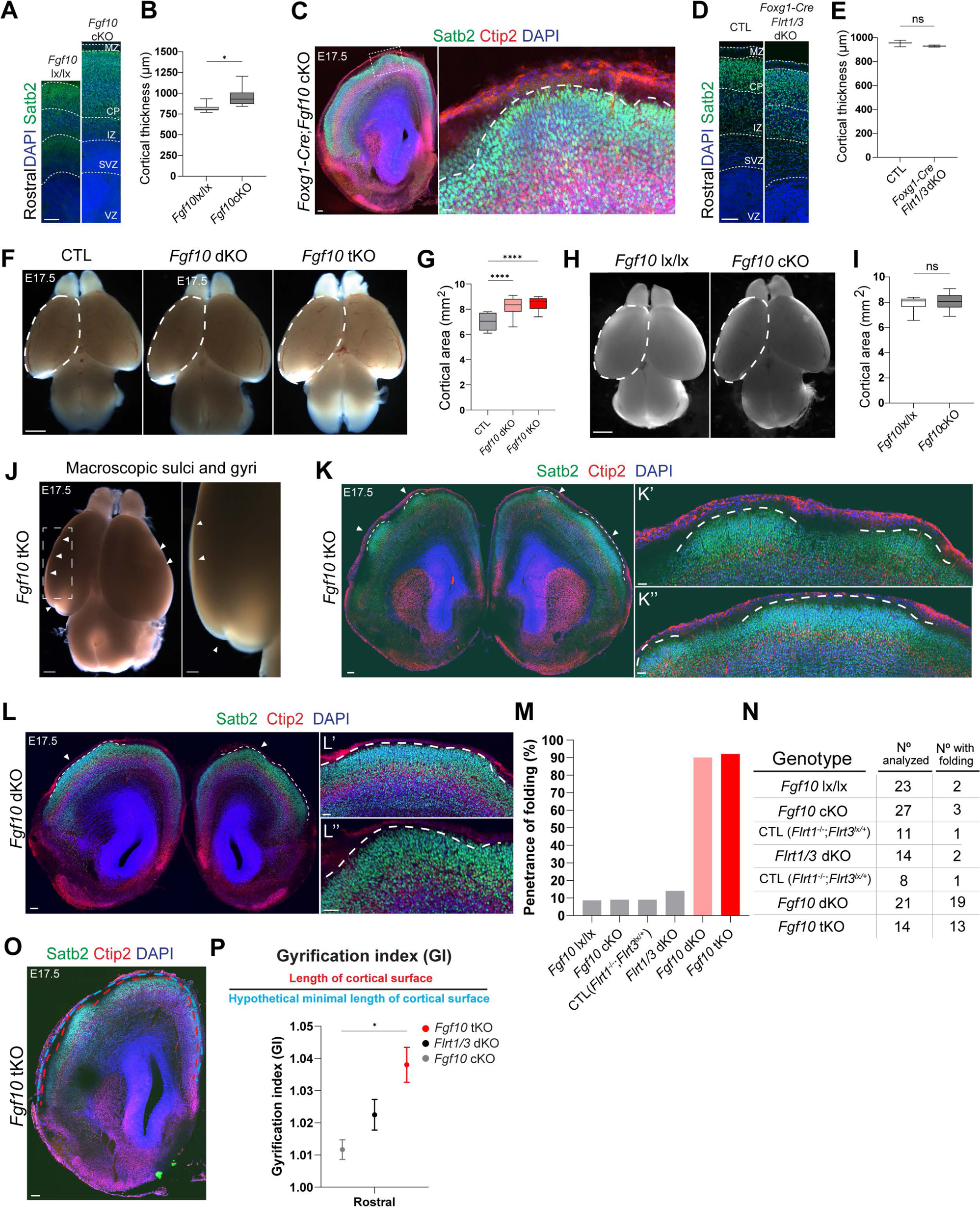
Loss of Fgf10 and Flrt1/Flrt3 enhances cortical folding and the presence of gyri. (A) Representative Satb2 (green) and DAPI (blue) stained E17.5 rostral sections of *Fgf10* lx/lx control and *Fgf10* cKO mice. (B) Quantifications of cortical laminar thickness shown in (A). Measurements were obtained from the bottom of the apical surface of the VZ to the pial surface. *Fgf10* lx/lx, n = 6 brains; *Fgf10* cKO, n = 6 cortices from 3 brains, p = 0.033, *p < 0.05, t-test with Welch correction. (C) Representative E17.5 Foxg1-Cre;*Fgf10* cKO brain section labeled with Satb2 (green), Ctip2 (red), and DAPI (blue). Area in dashed rectangle is shown with higher magnification on the right. Dashed line indicate a gyrus. (D) Representative Satb2 (green) and DAPI (blue) stained rostral sections of *Flrt1*^−/−^*;Flrt3* ^lx/+^ (CTL) and *Flrt1/3* dKO mice. (E) Quantifications of cortical laminar thickness shown in (C). Measurements were obtained from bottom of the apical surface of the VZ to the pial surface. CTL (*Flrt1*^−/−^*;Flrt3* ^lx/+^), n = 3 brains; *Flrt1/3* dKO, n = 3 brains. p = 0.282, t-test with Welch correction. (F) Representative whole-mount images of E17.5 CTL (*Flrt1*^−/−^*;Flrt3* ^lx/+^), *Fgf10* dKO and *Fgf10* tKO brains. Dashed areas were measured to obtain quantifications in (F). (E) Quantifications of the cortical areas shown in (E). CTL (*Flrt1*^−/−^*;Flrt3* ^lx/+^), n = 8 brains; *Fgf10* dKO, n = 22 brains; *Fgf10* tKO n = 13 brains. ****p < 0.0001, one-way ANOVA, Tukey Post hoc test. (H) Representative whole-mount images of E17.5 *Fgf10* lx/lx and *Fgf10* cKO brains. Dashed areas were measured to obtain quantifications. (I) Quantifications of the cortical areas shown in (G). *Fgf10* lx/lx, n = 21 brains; *Fgf10* cKO, n = 24 brains. p = 0.393, unpaired t-test with Welch correction. (J) Macroscopic sulci and gyri in E17.5 *Fgf10* tKO embryo. The area in the dashed rectangle is shown with higher magnification on the right, sulci and gyri are indicated by arrowheads. (K-K’’) E17.5 *Fgf10* tKO brain section stained with Satb2, Ctip2, and DAPI. Dashed lines indicate bilateral cortical folds, gyri. Higher magnifications on the right (K’ and K’’). (L-L’’) E17.5 *Fgf10* dKO brain section stained with Satb2, Ctip2, and DAPI. Dashed lines indicate bilateral gyri. Higher magnifications on the right (L’ and L’’). (M) Folding penetrance of E17.5 *Fgf10* lx/lx, *Fgf10* cKO, CTL (Flrt1^−/−^;Flrt3^lx/+^), *Flrt1/3* dKO, *Fgf10* dKO, and *Fgf10* tKO. All folds were located on the lateral side of the cortex. (N) Table of cortical folding penetrance at E17.5 for the indicated genotypes. Brains were analyzed for the presence of one or more sulci as shown in Figure 3M. (O) Representative immunostained image of a *Fgf10* tKO embryo depicting the quantification of the Gyrification Index (GI). Red line indicates the de facto length of the cortical surface; blue lineindicates the hypothetical minimal length of the cortical surface. (P) Quantification of GI values in E17.5 embryos. n= 6 brains *Fgf10* cKO (gray), n= 4 brains *Flrt1/3* dKO (black) and n=15 sections from a total of 7 brains *Fgf10* tKO (red) in the rostral region of brain. Data are shown as mean ± SEM; p = 0.014, *p < 0.05, one-way ANOVA with Tukey’s post hoc analysis. Scale bars represent 100 μm.(A), 200 μm, 200 μm (C), 100 μm (D), 1 mm (F), 1 mm (H), 500 μm, 500 μm (J), 100 μm, 50 μm, 50 μm (K), 100 μm, 50 μm, 50 μm (L), and 100 μm (O).

### Increased numbers of apical progenitors in *Fgf10* tKO mice

Ablation of Fgf10 was previously shown to expand apical progenitors, eventually resulting in an overproduction of neurons in the rostral cortex^25^. To investigate the underlying mechanism of the enhanced folding phenotype in *Fgf10* tKO mice, we performed quantifications of apical progenitors (Sox2), IPs (Tbr2) and mitotic dividing cells (pH3) in rostral cortical sections of E11.5 *Fgf10* tKO embryos. As expected, we found that Sox2-positive apical progenitors were expanded in *Fgf10* dKO and tKO mice compared to littermate controls (*Flrt1*^−/−^;*Flrt3*^lx/+^) (Figure 4 A,B). Unlike *Cep83* tKO mice, the numbers of Tbr2-positive IPs and double positive PH3/Pvim cells were not increased in E11.5 or E13.5 *Fgf10* dKO and tKO embryos (Figure 4 A,B; Suppl. Figure S4 A,B,D,E). At E11.5, the proportions of mitotic cells (pH3+) between apical versus basal layers were shifted towards apical layers in *Fgf10* dKO and tKO embryos (Figure 4C), suggesting increases in dividing NECs and aRG cells. A somewhat weaker effect could still be seen in E13.5 embryos consistent with Fgf10’s described early and transient effects^25^ (Suppl. Figure S4 C). Collectively, these results suggest that deletion of Fgf10/Flrt1/3 substantially increases the division of early apical progenitors, NECs and aRG cells, which magnifies the cortex folding phenotype seen in the *Flrt1/3* dKO mice. Together with the findings in *Cep83* tKO mice, these results indicate that expansion of cortical progenitors in conjunction with alteration of cell migration drives folding of the mouse cortex. The types of folds (sulci versus gyri) appear to depend on the types of progenitors that were expanded: expansion of apical progenitors in *Fgf10* tKO mice correlated with the appearance of gyri, while expansion of IPs and bRG cells in *Cep83* tKO mice correlated with the appearance of sulci.

**Figure 4.**
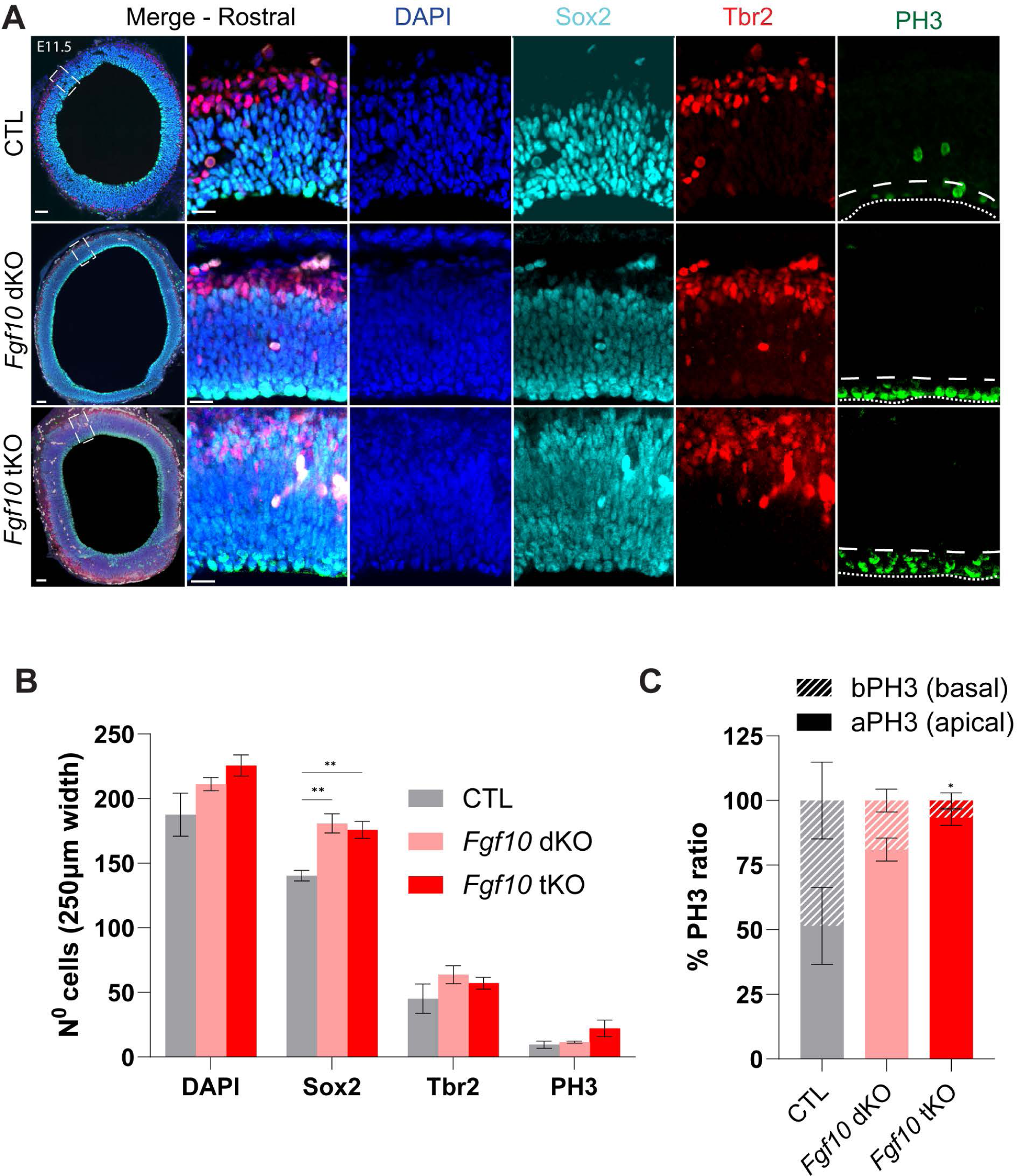
Progenitor expansion in *Fgf10* tKO *and Fgf10* dKO embryos at E11.5. (A) E11.5 rostral cortices of CTL (*Flrt1*^−/−^*;Flrt3* ^lx/+^), *Fgf10* dKO and *Fgf10* tKO embryos were stained DAPI (blue), apical progenitors Sox2 (cyan), intermediate progenitors Tbr2 (red), and mitotic cells PH3 (green). Apical and basal sides of the VZ are indicated with dotty and dashed lines in PH3 stained images, respectively. Areas in dashed rectangles in (A) are shown with higher magnification on the right. (B) Quantification of the data shown in (A). CTL (*Flrt1*^−/−^*;Flrt3* ^lx/+^) n = 5 brains, *Fgf10* dKO n = 5 brains, *Fgf10* tKO n = 5 brains. Data are shown as mean ± SEM; CTL vs. *Fgf10* dKO p = 0.002, CTL vs. *Fgf10* tKO p = 0.006, *Fgf10* dKO vs. *Fgf10* tKO p = 0.810. **p < 0.01, one-way ANOVA with Tukey’s post hoc analysis. (C) Proportion of apical/basal mitotic cells (PH3) in rostral region CTL (*Flrt1*^−/−^*;Flrt3* ^lx/+^) n = 5 brains, *Fgf10* dKO n = 5 brains, *Fgf10* tKO n = 5 brains. Data are shown as mean ± SEM; CTL vs. *Fgf10* dKO p = 0.097, CTL vs. *Fgf10* tKO p = 0.018, *Fgf10* dKO vs. *Fgf10* tKO p = 0.617. *p < 0.05, one-way ANOVA with Tukey’s post hoc analysis. Scale bars represent 50 μm, 20 μm (A).

### Higher cell density in gyri of *Fgf10* tKO mice

To further characterize the cortical folds in both tKO mouse models, we quantified and compared the cell densities within the folded areas. Naturally forming gyri in gyrencephalic species tend to have a higher cell density than sulcus areas^3,7,40,41^. It was therefore interesting to ask, if the generated gyri in *Fgf10* tKO mice would have higher cell densities than control areas in the same cortical region. We quantified the densities of Satb2-positive upper-layer cortical neurons and found that the cell densities in the gyrus regions were approximately 50% higher than those in adjacent areas or in an equivalent region of a littermate control brain (Figure 5 A-C). In contrast, the densities of Satb2-positive neurons in the sulcus regions of *Cep83* tKO mice were on average similar to those in other regions of the same section or control brains (Figure 5 D-F). These findings confirm that the cortical folds of *Fgf10* tKO mice display features of naturally occurring gyri.

**Figure 5.**
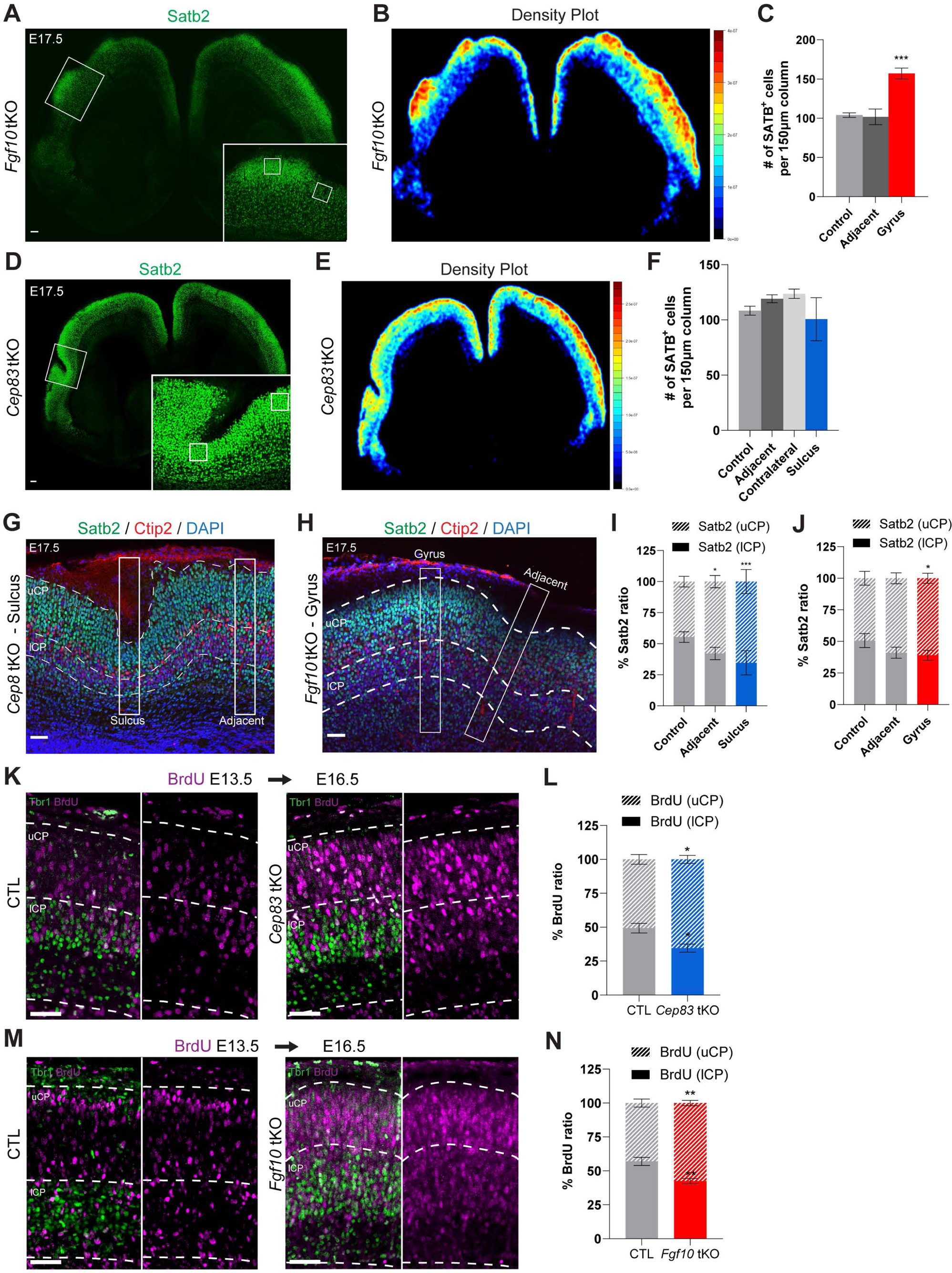
Higher cell density in gyri of *Fgf10* tKO mice. (A) Coronal brain sections of *Fgf10* tKO at E17.5, immunostained with Satb2, upper layer CP neuronal marker. Area in dashed rectangle is shown with higher magnification in the lower right corner. Small squares depict the ROIs used for quantification. (B) Heatmap of Satb2+ cell density of the section shown in (A). The color bar indicates a higher (red) or lower (blue) density of Satb+ cells. (C) Quantification of the data shown in (A). Satb2+ cells were counted in gyri and adjacent region of the cortex and in littermate controls (n = 5 brains). Data are shown as mean ± SEM; Control vs. Adjacent p = 0.975, Control vs. Gyrus p = 0.001, Adjacent vs. Gyrus p = 0.0004. ***p < 0.001, one-way ANOVA with Tukey’s post hoc analysis. (D) Coronal brain sections of *Cep83* tKO at E17.5 immunostained with Satb2. Area in dashed rectangle is shown with higher magnification in the lower right corner. Small squares depict the ROIs used for quantification. (E) Heatmap of Satb2+ cell density of the section shown in (D). The color bar indicates a higher (red) or lower (blue) density of Satb+ cells. (F) Quantification of the data shown in (D). Satb2+ cells were counted in sulci and adjacent and contralateral side of cortices and littermate controls. (n = 6 brains). Data are shown as mean ± SEM; Control vs. Adjacent p = 0.879, Control vs. Contralateral p = 0.723, Control vs. Sulcus p = 0.952, Adjacent vs. Contralateral p = 0.990, Adjacent vs. Sulcus p = 0.594, Contralateral vs. Sulcus p = 0.415. one-way ANOVA with Tukey’s post hoc analysis. (G) Sulcus and adjacent region from an E17.5 *Cep83* tKO section immunostained with Satb2 (green, upper CP) and Ctip2 (red, lower CP), DAPI (blue). White rectangles depict the ROIs used for quantifications shown in I. (H) Gyrus and adjacent region from an E17.5 *Fgf10* tKO section immunostained with Satb2 (green), Ctip2 (red), and DAPI (blue). White rectangles depict the ROIs used for quantifications shown in J. (I) Quantification of data shown in (G). Proportion of Satb2+ cells in the upper and lower CP (defined by Ctip2 staining) in sulcus region, versus areas adjacent to the sulcus, and similar ROIs in CTL mice (*Flrt1*^−/−^;*Flrt3*^lx/+^) (n = 6 brains per group). Data are shown as mean ± SEM; lCP: Control vs Adjacent p = 0.012, Control vs. Sulcus p = 0.0002, Adjacent vs. Sulcus p = 0.131. uCP: Control vs. Adjacent p = 0.011, Control vs. Sulcus p = 0.0003, Adjacent vs. Sulcus p = 0.180 *p <0.05, ***p < 0.001, one-way ANOVA with Tukey’s post hoc analysis. (J) Quantification of data shown in (H). Details see panel (I) (n = 3 brains per group). Data are shown as mean ± SEM; Control vs. Adjacent p = 0.089, Control vs. Gyrus p = 0.041, Adjacent vs. Gyrus p = 0.810. *p < 0.05, one-way ANOVA with Tukey’s post hoc analysis. (K) BrdU injection was performed at E13.5 and analyzed at E16.5 in CTL (*Flrt1*^−/−^*;Flrt3* ^lx/+^)and *Cep83* tKO embryos. BrdU expression was confirmed by immunostaining coronal sections with BrdU (magenta). The CP was subdivided into uCP and lCP using Tbr1 (green), and the number of BrdU+ neurons in each layer was quantified in (L). (L) Quantification of the data shown (K). n = 4 embryos, Data are shown as mean ± SEM; p = 0.017, *p < 0.05, t-test with Welch correction. (M) BrdU injection and analysis in CTL (Flrt1−/−;Flrt3lx/+) and *Fgf10* tKO embryos as described in panel K. (N) Quantification of the data shown in (M). n = 5 sections from 4 embryos, Data are shown as mean ± SEM; p = 0.005, **p < 0.01, t-test with Welch correction. Scale bars represent 100 μm (A), 100 μm (D), 50 μm (G), 50 μm (H), 50 μm (K), and 50 μm (M)

### Increased neuronal migration speed in *Cep83* tKO and *Fgf10* tKO mice

We had previously reported that loss of Flrt1/3 resulted in cortical layer thinning in sulcus areas, particularly in lower CP, similar to gyrencephalic species such as the ferret^42^ and primates^43^. We found a similar reduction in the thickness of deeper layers in the folded areas of *Cep83* tKO mice, but not in the *Fgf10* tKO mice, compared to controls (*Flrt1*^−/−^*Flrt3*^lx/+^) (Figure 5 G,H). This is consistent with the *Cep83* tKO favoring sulcus and the *Fgf10* tKO favoring gyrus formation. The proportion of Satb2+ cells in upper and lower CP shifted towards upper layer CP in both mutants (Figure 5 I,J) consistent with previous observations after ablation of Flrt1/3^31^ and with mutant pyramidal neurons migrating faster through the cortical plate. Support for such a model came from bromodeoxyuridine (BrdU) pulse labeling of newborn pyramidal neurons at E13.5 and analyzing their distribution in the CP at E16.5. In both tKO models, we observed a shift in the proportion of BrdU+ neurons in the upper versus lower CP compared to their respective controls, and this shift occurred throughout the CP independent of folded areas (Figure 5 K-N).

Previously, we had used ex vivo live imaging of embryonic cortices to demonstrate that loss of Flrt1/3 increases neuronal migration speed^31^. We took the same approach to analyze the speed of radial migration in both tKO models, although these experiments were extremely challenging due to the low yield of *in utero* electroporation (IUE) of tKO embryos. IUE with pCAG-Ires-GFP was performed at E13.5 and time-lapse imaging was done for up to 24h from E15.5 cortical slices. Migrating GFP-labelled neurons were tracked within the CP and the average speeds were calculated. The results suggested that the average speeds of *Cep83* tKO and *Fgf10* tKO neurons were faster than control neurons (Suppl. Figure S5). Together these results suggest that the enhanced cortical folding phenotype in the triple KO mice is the result of combining progenitor expansion with faster migration.

### Modeling cortex folding in *Cep83* tKO and *Fgf10* tKO mice

Previously, we had used data-driven computational simulations to provide evidence that cell clustering and increased migration speed produced cortical folds^31^. To model the combined effects of progenitor expansion and cell migration on cortex folding in the tKO mice, we first analyzed the global cellular changes in the tKO mice by scRNAseq. We dissected the rostral and medial parts of the neocortex from three E15.5 *Cep83* tKO and three *Fgf10* tKO embryos plus respective controls and performed scRNAseq to identify the different cortical cell types (Figure 6A). We obtained 19268 and 14777 single cell transcriptomes from *Cep83* tKO and *Fgf10* tKO embryos, respectively (with a median of 3998 and 3338 genes per cell), performed principle component analysis on the scaled gene expression data, followed by UMAP visualization and unsupervised clustering. We assigned clusters to different cell types based on the co-expression of multiple marker genes (Figure 6 B,E; Suppl. Figure S6 A-C). In support of our immunostaining results, *Cep83* tKO embryos had an expanded Eomes/Tbr2-positive IP pool compared to control embryos, while the proportions of other cell types remained largely unaffected (Figure 6 C,D). *Fgf10* tKO embryos did not show an expanded apical progenitor pool, presumably because the window of FGF10-mediated differentiation of apical progenitors was much earlier in development^25^ (Suppl. Figure S6 D,E). *Cep83* tKO embryos also showed an increased proportion of upper layer neurons consistent with the expanded pool of IPs (Figure 6 E-G), an effect that was not seen in *Fgf10* tKO embryos (Suppl. Figure S6 F-G). Given that FLRT-expressing and non-expressing migrating neurons have different cell adhesion properties that influence their migration behavior^31^, we quantified their proportions. We found that the numbers of Flrt3-mutant neurons (previously destined to express Flrt3), were increased compared to Flrt3-negative neurons in *Cep83* tKO embryos (60:40 proportion), and decreased in *Fgf10* tKO embryos (45:55 proportion) (Figure 6 H,I). Moreover, the overall increases in cell numbers in the two tKO models varied considerably, with 9% in E13.5 *Cep83* tKO compared to 33% in E13.5 *Fgf10* tKO embryos (Figure 6 FJ).

**Figure 6.**
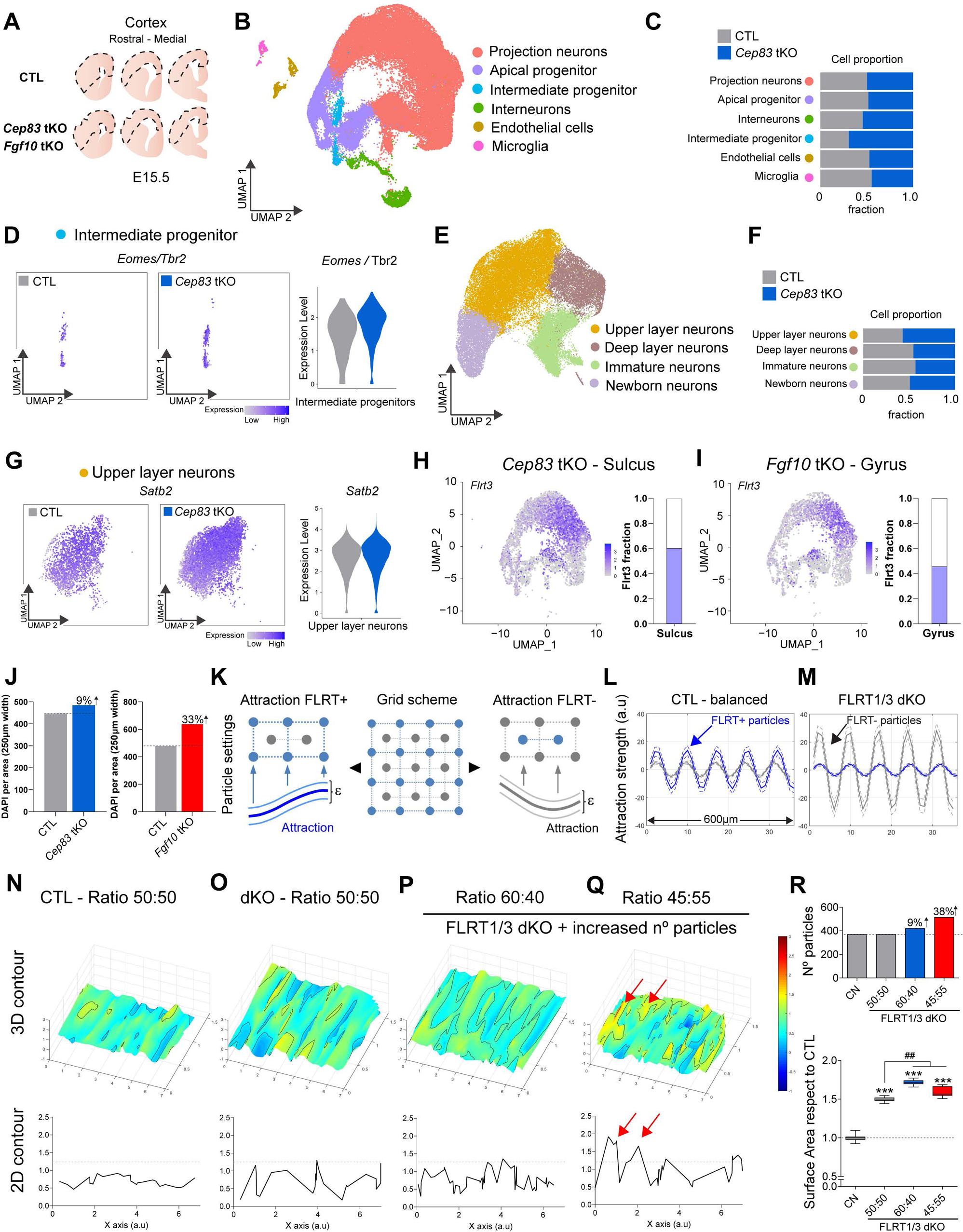
Modeling cortex folding in *Cep83* tKO and *Fgf10* tKO mice. (A) Scheme of tissue collection for scRNAseq. Areas indicated by dashed line were manually dissected from coronal sections of E15.5 rostral to medial cortex. Three *Cep83* tKO brains (one with sulcus), three *Fgf10* tKO brains (one with gyrus), n=2 littermate CTLs each (*Flrt1*^−/−^*;Flrt3* ^lx/+^) were collected for tissue preparation. (B) UMAP clustering of cells from E15.5 *Cep83* tKO (n = 19267), *Fgf10* tKO (n = 14777), CTL (*Flrt1*^−/−^*;Flrt3* ^lx/+^) (n = 22107) embryos classified as projection neurons (n = 40900), apical progenitors (n = 9212), interneurons (n = 2778), intermediate progenitors (n = 1474), endothelial cells (n = 1357), and microglia (n = 531). (C) Proportions of each cell type by genotype in CTL (*Flrt1*^−/−^*;Flrt3* ^lx/+^) and *Cep83* tKO mice. (D) Expression of the IP marker gene Eomes (Tbr2) in intermediate progenitor cell type for CTL (*Flrt1*^−/−^*;Flrt3* ^lx/+^) and *Cep83* tKO embryos. (UMAP and violin plot). (E) UMAP clustering of projection neurons with annotated subpopulations, upper layer neurons (n = 17963), immature neurons (n = 10103), deep layer neurons (n = 6897), and newborn neurons (n = 5937). (F) Proportions of subpopulations by genotype in CTL (*Flrt1*^−/−^*;Flrt3* ^lx/+^) and *Cep83* tKO embryos. (G) Expression of CP layer marker gene, Satb2, in upper layer neurons for CTL (*Flrt1*^−/−^*;Flrt3* ^lx/+^) and *Cep83* tKO embryos. (UMAP and violin plots). (H) UMAP clustering and quantification of Flrt3+ cells in apical progenitors, intermediate progenitors, and projection neurons for *Cep83* tKO embryo in sulcus area. (I) UMAP clustering and quantification of Flrt3+ cells in apical progenitors, intermediate progenitors, and projection neurons for *Fgf10* tKO embryo in gyrus area. (J) Number of cells based on DAPI staining per area in CTL (*Flrt1*^−/−^*;Flrt3* ^lx/+^) and *Cep83* tKO and *Fgf10* tKO (K) Scheme illustrating the position of the FLRT-positive (blue) and –negative (grey) particles. Lines between the particles indicate attraction forces that are random within a small range (ε). (L,M) Sinus equations representing the attraction forces among FLRT-positive (blue) and negative (grey) particles in both control (L) and FLRT1/3 dKO condition (M). In the latter, FLRT-negative particles show increased adhesion forces (higher amplitude) than FLRT-positive ones. Dashed lines indicate the small range of variation (ε). (N-Q) Surface plots and line graphs of the distribution of the particles on the Z axis after computer simulations. The control situation with a 50:50 ratio of FLRT+ and FLRT-negative particles results in a uniform surface (N). The Flrt1/3 dKO condition with altered attraction forces results in a wavy surface (O). This effect is enhanced by increasing the number of particles and changing the proportions between FLRT-positive and -negative particles (P,Q). 2D plots representing the contour of a central slice through the surface plot are shown below. Red arrows indicate elevated areas that are more prominent in the Q condition. (R) Number of particles and surface area calculated from conditions N-Q. Data are shown as mean ± SEM; ***p < 0.001 vs Control and ## p < 0.01 vs 50:50, one-way ANOVA with Tukey’s post hoc analysis.

Similar to our previous modeling analysis^31^, we distributed particles representing Flrt1/3-positive (blue) and -negative cells (grey) in a 2D grid in a 50:50 ratio (Figure 6 K). The adhesive forces between cells were modeled as sine equations, reflecting the repeated patterns of cell clusters in Flrt3 gain-and loss-of-function experiments^44,31^ (Figure 6 L,M). In the wild-type (balanced) situation, we determined that Flrt1/3-positive particles were attracted to one another (via FLRT-mediated homophilic adhesion) and the attraction force was higher than the one between Flrt1/3-negative cells (Figure 6 L). In the *Flrt1/3* dKO situation, we defined that Flrt1/3-mutant particles showed reduced attraction to one another (due to the loss of FLRT-mediated homophilic adhesion) (Figure 6 M). We previously assumed that they were segregated due to repulsion from Flrt1/3-negative particles^31^. However, more recent analyses revealed that instead the segregation occurred because FLRT1/3-negative cells gained increased adhesion toward each other and that the adhesive force was stronger than homophilic adhesion between FLRT1/3 mutant cells (Y. Shen, unpublished observations). These results suggest a model in which FLRT1/3 mutant cells are sorted by differential adhesion, consistent with the hypothesis of cell segregation by differences in intercellular adhesiveness^45^. We incorporated this new result into our model for *Flrt1/3* dKO and tKO mice, adjusting the amplitudes of the sine equations to increase the strength of the intercellular adhesion among FLRT1/3-negative cells in comparison to FLRT1/3 mutant cells (Figure 6 L,M). To analyze the behavior of the moving particles, they were set to move along the z axis, and both speed and attraction forces were random within a small range (ε) to resemble the fluctuations present in biological systems^46^ (Figure 6 K). In the balanced situation, with a slightly higher attraction between FLRT1/3-positive particles compared to FLRT1/3-negative particles, the particles formed a rather smooth surface after moving along the z axis (Figure 6 N). In the *Flrt1/3* dKO situation with low attraction between FLRT1/3-mutant cells and strong attraction between FLRT1/3-negative cells, but no change in cell proportions or density, the surface became more wrinkled, consistent with the cortex folding phenotype (Figure 6 O,R). In the *Cep83* tKO situation with similar changes in attraction forces as in *Flrt1/3* dKO mice, a 60:40 proportion of FLRT1/3-mutant to negative particles, and an overall increase of 9% cell density, the surface became even more wrinkled, consistent with the enhanced cortex folding phenotype featuring mostly sulci (Figure 6 P,R). Conversely, the *Fgf10* tKO situation, with a 45:55 Flrt1/3-mutant to negative ratio and a 33% increase in cell density, resulted in a surface that had wrinkles, but also elevated areas, resembling the enhanced cortex folding phenotype featuring predominantly gyri (Figure 6 Q,R). Taken together, the combination of modeling attraction forces, proportions of FLRT1/3-mutant to FLRT1/3-negative particles, and cell densities matched the experimental observations with dKO and tKO mice rather well by producing a more wrinkled surface with sulcus-like and gyrus-like features.

## DISCUSSION

In the present study, the genetic combination of progenitor expansion with divergent migration allowed us to address a long-lasting question concerning the mechanisms of cortex folding: what is the relative contribution of progenitor expansion and cortical migration to this complex process? Previously, we have shown that genetic deletion of Flrt1/3 adhesion molecules causes formation of cortical sulci resulting from increased lateral dispersion and faster neuron migration, in the absence of progenitor amplification^31^. Here, we find that combining the *Flrt1/3* dKO migration model with an additional genetic deletion that causes progenitor expansion greatly enhances cortex folding. The main features of cortex folding, gyri versus sulci, correlate with the type of progenitor pool that was unevenly enlarged. Expansion of intermediate progenitors by deletion of Cep83 results in a magnified cortex folding phenotype with the presence of sulci. Enlargement of the apical progenitor pool by deletion of Fgf10 results in enhanced cortex folding by promoting gyri formation. Single cell transcriptomics and computational simulations identify key parameters that cooperate to promote cortical gyrification: changes in adhesive properties of cortical neurons, their proportions and densities in the cortical plate, combined with lateral dispersion during their radial migration.

### Mechanisms underlying cortical gyrification

Most previous models of cortex folding involved the expansion of progenitors^10,11,18,47^. The *Flrt1/3* dKO mouse model was therefore quite unique in the sense that cortex folding was induced independent of progenitor amplification, by lateral dispersion and accelerated neuron migration^31^. This neuronal migration behavior remained an important feature of the novel tKO models used in the present study and apparently was affected neither by deleting Cep83 nor Fgf10. Notably, both tKO models combined the divergent migration with progenitor expansion. And yet, the folding phenotypes of the two tKO models were qualitatively very different. What could be the critical parameters determining the formation of sulci versus gyri?

In *Cep83* tKO embryos, where sulci are prominent, progenitor expansion starts around E12.5, involving IPs throughout the entire cortex. scRNAseq analysis revealed a higher proportion of FLRT3-mutant (previously destined to co-express FLRT1/3) to FLRT3-negative neurons (60:40) than in controls (50:50), accompanied by a modest 9% increase in cell density. Conversely, in *Fgf10* tKOs which feature predominantly gyri, progenitor expansion happens early, involves NECs and apical progenitors, and is confined to the rostral parts of the cortex. scRNAseq analysis revealed a reduced proportion of FLRT3-mutant to FLRT3-negative neurons (45:55), coupled with a strong 33% increase in cell density.

Which of these changes may be most important? We have previously shown that sulcus formation in the *Flrt1/3* dKO model is highest in the lateral cortex where both proteins are strongly expressed^31,44^. In the *Cep83* tKO model, more FLRT-mutant cells are generated, and the phenotype extends into the lateral cortex, inducing cortical expansion in an area where divergent and faster neuronal migration is prominent due to the absence of FLRT1/3 proteins^31^. This may favor the strengthening of the phenotype already present in the *Flrt1/3* dKO, namely the formation of sulci. Conversely, in the *Fgf10* tKO model, fewer FLRT3-mutant cells are generated compared to the *Flrt1/3* dKO model, weakening lateral dispersion and prompting neurons to taking a more radial migratory path. In addition, the effects of Fgf10 deletion are confined to the rostral cortex where the expression of FLRT3 protein is low^31^. This local overproduction of predominantly radially (rather than tangentially) migrating neurons may favor the formation of gyri (see model in Figure S6 H). Moreover, it is possible that differences in cell densities between the two models influence the formation of sulci versus gyri. Deletion of Cep83 resulted in a modest increase in cell density and the formation of sulci, whereas deletion of Fgf10 featured a large increase in cell density and the formation of gyri. This would be consistent with studies showing that in gyrencephalic species, regions in the cortical plate with lower and higher neuronal densities correspond to future sulci and gyri, respectively, during development^48^.

Our model is also supported by computational simulations. Simply reducing intercellular adhesion between FLRT-mutant particles (as in the *Flrt1/3* dKO model) and increasing intercellular adhesion between FLRT-negative particles (Y. Shen, unpublished observations), while keeping particle numbers unchanged, generated a more wrinkled surface. We propose a modification of our previous model^31^ that FLRT1/3 mutant cells are sorted away from FLRT-negative cells by differential adhesion (rather than being subjected to increased repulsion from FLRT-negative cells) and that this effect favors lateral dispersion and faster migration. The folding (wrinkling) effect was enhanced when more FLRT-mutant particles were added and the particle density was modestly increased as in the *Cep83* tKO model. In this case, the combination of more particles with reduced intercellular adhesiveness with a modest cell density increase favored sulci formation. The wrinkled surface contained elevated areas when fewer than normal FLRT-mutant particles were added and the particle density was markedly increased as in the *Fgf10* tKO model. In that case, the combination of more particles with increased intercellular adhesiveness with a large increase in cell density favored the formation of gyri.

Our findings provide a conceptual framework of how the gyrencephalic cortex structure is physically sculpted during development. It incorporates the roles of progenitor expansions and the effects these expansions have on overall cell density and on the proportions of cortical neurons with distinct adhesive and migratory properties. In the future, it would be interesting to test other combinations of genetic manipulations that expand different types of progenitors and lead to divergent neuron migration. Since the human gyrencephalic neocortex expresses much lower levels of FLRT1/3 compared to the lissencephalic mouse neocortex^31^, these studies will also provide insights into the transition from gyrencephaly to lissencephaly during mammalian evolution.

## ACKNOWLEDGEMENTS

We thank S. Cappello for her insightful comments during the course of this study, C. Mayer, A. Bright, Y. Kotlyarenko, P.A. Morales and H. Lim for help with scRNA sequencing experiment, K. Voelkl and L. Schaffmayer for technical help with scRNA sequencing experiment. D. Feigenbutz for help with data processing of scRNA library. This study was supported by the Max-Planck Society, by the Ministry of Science and Technology of China (2021ZD0202300) and New Cornerstone Investigator Program (S.-H.S.). by the Deutsche Forschungsgemeinschaft (DFG, German Research Foundation) under Germany’s Excellence Strategy within the framework of the Munich Cluster for Systems Neurology (EXC 2145 SyNergy – ID 390857198).

## AUTHOR CONTRIBUTIONS

S.C. and R.K. conceptualized the study and designed experiments; S.C. performed most of the experiments; D.S.D helped with phenotypic analysis of some of the mutant mice and time-lapse experiment; M.T and A.E. performed 3D imaging; T.S. performed scRNAseq data analysis; T.R. performed time-lapse speed analysis; G.S.B helped with ISH and immunostainings; Y.S. assisted in scRNAseq experiment; D.d.T. performed computational simulations; W.C., J.Y., and S.-H.S. contributed reagents for the paper; R.K and D.d.T. supervised; R.K., S.C, and D.d.T wrote the manuscript; R.K. provided funding.

## STAR+METHODS

### Mouse lines

Flrt1^−/−^; Flrt3^lx/lacZ^ mice^49,50^ were crossed with either Cep83^lx/lx^ mice^18^ or Fgf10^lx/lx^ mice^51^, leading to *Cep83^lx/lx^;Flrt1^−/−^; Flrt3^lx/lacZ^* mice or Fgf10^lx/lx^;Flrt1^−/−^; Flrt3^lx/lacZ^ mice. These animals were further crossed with either Emx1-Cre or Foxg1-Cre to remove the floxed alleles. All animal experiments were approved by the Government of Upper Bavaria and carried out in accordance with German guidelines for animal welfare. All mice (C57BL/6 and 129/SvJ mixed background) were housed with 12:12h light/dark cycle and food/water available *ad libitum* in the facilities of the Max Planck Institute of Biological Intelligence. With regard to embryonic stage, the midday of vaginal plug formation was regarded as embryonic day 0.5 (E0.5).

### Immunohistochemistry

For tissue preparation, whole heads from E11.5, E13.5 mice and whole brains from E15,5, E17,5 mice were dissected and were fixed in 4% PFA in PBS at 4 °C overnight and stored in PBS. To obtain coronal brain sections, tissue was embedded in 4% agarose in PBS. Serial 100 µm (E15.5 and E17.5) and 80 µm (E11.5 and E13.5) thick sections were cut in PBS with a Leica VT1000 S vibratome. The sections were incubated with primary antibodies diluted with 0.2% BSA/0.5%Triton/5% donkey serum in PBS, including rabbit anti-SATB2 (1:500, Abcam), rat anti-Ctip2 (1:500, Abcam), goat anti-SOX2 (1:1000, R&D Systems), rat anti-Histone H3 (1:500, Abcam), rabbit anti-Tbr1 (1:500, Abcam), rabbit anti-Tbr2/Eomes (1:200, Abcam), rabbit anti-Cux1 (1:500, Santa Cruz), mouse anti-Pvim (1:500, Abcam) and rat anti-BrdU (1:100, Abcam). Secondary antibodies were diluted at 1:500 with 0.2% BSA/0.5%Triton/5% donkey serum in PBS and incubated for 2 hr at room temperature. The secondary antibodies used were Cy3-conjugated donkey anti-rat IgG, Cy3-conjugated donkey anti-rabbit IgG, Cy3-conjugated donkey anti-mouse IgG, Alexa 488-conjugated donkey anti-rabbit IgG, Alexa 488-conjugated donkey anti-rat IgG, and Alexa 674-conjugated donkey anti-goat IgG (Jackson Immunoresearch Laboratories). Cell nuclei were counterstained with 4’,6-Diamidino-2-phenylindole dihydrochloride (DAPI)/PBS (1:1000; Invitrogen). For observation, sections were mounted with DAKO mounting medium, and sealed with cover slides by Menzel-Gläser (Menzel GmbH & CO). Images were visualized using a Leica SP8 confocal microscope.

### *In utero* electroporation assays

*In utero* electroporation was performed as described previously with minor modifications (Saito and Nakatsuji, 2001, Sato et al., 2013). Timed pregnant mutant mice at E13.5 were anesthetized with isoflurane (CP-pharma) using a Fluovac 34-0387 (Harvard Apparatus) and Vevo Compact Anesthesia System (VisualSonic). For surgery, the uterus of the mouse was exposed after making an approximately 3 cm incision in the middle abdominal region. Then, both sides of uterine horns were pulled out, and approximately 1-2 μl of a mixture containing 1 μg/μl pCAG-Ires-GFP plasmid (Addgene, Cat #11159) and 1% Fast Green (Sigma, final concentration 0.2%) in PBS was injected into the lateral ventricle of embryos with a mouth-controlled glass capillary pipette. Immediately, square pulses (30 V, 50 ms, six times at 1 s intervals) were delivered into embryos with an electroporator (ECM 830, BTX) and a forceps-type electrode (CUY650P5, NepaGene). After the electroporation, the uterus was returned back inside the abdomen using ring forceps and the incision was sutured with PERMA-HAND Seide (9.3 mm diameter curved needle, 45 cm of thread, Ethicon). After surgery, mice were placed on a 37°C heating pad for recovery and kept until the desired embryonic stage.

### BrdU analysis

A single injection of 5-bromo-20-deoxyuridine (BrdU, Sigma-Aldrich) dissolved in PBS, 0.15-0.2 ml at a concentration of 10 mg/ml BrdU was administrated to pregnant females at E13.5 and E15.5 intraperitoneally for a short BrdU pulse analysis, to give a final concentration of 50 mg per g of mouse weight. Pregnant females were then sacrificed after 30 mins. Brains were collected and fixed overnight with 4% PFA and stored in PBS. For staining with the BrdU antibody, sections were pre-treated with 2N HCl for 30 min at RT and washed with Na2B4O7 (pH 8.5) twice for 15 min.

### Time-lapse experiments

Embryo cortices were electroporated at E13.5 with pCAG-Ires-GFP (Addgene Cat #11159). After 2 days, E15.5 of embryonic brains were isolated in an ice cold sterile filtered and aerated (95% O2/5% CO2) dissection medium (15.6 g/l DMEM/F12 (Sigma), 1.2g/l NaHCO3, 2.9 g/l glucose (Sigma), 1% (v/v) penicillin streptomycin (GIBCO)). Using 4% low melting agarose (Biozym), brains were embedded for cutting into 300 mm thick sections using a vibratome (Leica, VT1200S). Sections were immersed in a collagen mix (64% (v/v) cell matrix type I-A (Nitta Gelatin), 24% (v/v) 5x DMEM/F12, 12% (v/v) reconstitution buffer (200 mM HEPES, 50mM NaOH, 260mM NaHCO3) and transferred onto a cell culture insert (Millicell, PICMORG50). Sections were then incubated at 37°C for 10 min to solidify the collagen mixture. 1.5 mL slice medium (88% (v/v) dissection medium, 5% (v/v) horse serum, 5% (v/v) fetal calf serum, 2% (v/v) B27 supplement (GIBCO), 1% (v/v) N2 supplement (GIBCO)) was added to the dish surrounding the culture insert and incubated for minimum 30 min at 37°C. Before imaging, culture medium was added to the top of the sections to allow objective immersion. Imaging was performed using a 20x water immersion objective on a Leica SP8 confocal microscope system equipped with a temperature-controlled carbon dioxide incubation chamber set to 37°C, 95% humidity, and 5% CO2. Sequential images were acquired every 15 min for 24-48 hr. After imaging, slices were genotyped to identify triple knock outs and littermate controls. The neuronal movement was tracked using the software Fiji with plugin ‘Manual Tracking.’ Only neurons entering the CP were tracked. Tracking was stopped when cells reached the upper cortical plate Single cell track analysis and plotting was carried out using custom made python scripts.

### 3DiSCO tissue clearing

A modified clearing protocol was used based on Belle et al., 2017. Briefly, E17.5 embryo brains were fixed in 4% PFA overnight at 4oC and stored in PBS. First, whole brain immunostaining was performed before tissue clearing. Brains were incubated in blocking buffer for 24h at RT on a horizontal shaker. Then brains were stained with primary antibodies, rabbit anti-SATB2 (1:1000, Abcam) and rat anti-Ctip2 (1:1000, Abcam) over 7 days with blocking buffer containing saponin at 37°C on a shaker. Next, brains were washed for 1 hour 6 times in 15 ml falcon tubes containing blocking buffer. Then, brains were incubated at 37oC on the horizontal shaker over 2 days with secondary antibodies Alexa Fluor 647 and 594 (1:500, Jackson Immunoresearch Laboratories), which had been filtered with a 0.20 mm filter and diluted in blocking buffer with saponin 0.1%. The brains were washed for 1 hour 6 times in 15 ml falcon tubes filled with blocking buffer. Then, tissue clearing was performed. Briefly, the whole stained brains were immersed in 50% tetrahydrofurane (THF, Carl Roth) overnight, in 80% THF for 1 h, in 100% THF for 1 h, then in fresh 100% THF for another hr, then in 100% dichloromethane (DCM, Carl Roth) until the brains sank, and finally in 100% dibenzylether (DBE, Sigma), followed by another step of fresh 100% DBE. Then, cleared brains were imaged with a 4× objective lens (Olympus XLFLUOR 340) equipped with an immersion-corrected dipping cap mounted on a LaVision UltraII microscope coupled to a white-light laser module (NKT SuperK Extreme EXW-12). Images were taken with 16-bit depth and at a nominal resolution of 1.625 μm per voxel on the x and y axes. Brain structures were visualized with Alexa Fluor 594 (using a 580/25 nm excitation filter and a 625/30 nm emission filter) and Alexa Fluor 647 fluorescent dye (using a 640/40 nm excitation filter and a 690/50 nm emission filter) in sequential order. Laser power was set for each channel so as not to exceed the dynamic range of the Neo 5.5 sCMOS camera (Andor). For 12× imaging, we used a LaVision objective (12×/0.53 NA MI PLAN with an immersion-corrected dipping cap). Camera exposure time was set to 105 ms and 90 ms for the 4× and 12× imaging respectively. In the z dimension, images were taken in 5 μm and 2 μm steps, while using left- and right-sided illumination for the 4× and 12× imaging, respectively. Our nominal resolution was 1.625 µm × 1.625 µm × 5 µm and 0.602 µm × 1.602 µm × 2 µm for the x, y and z axes, with 4× and 12× objectives. The thinnest point of the light sheet was 28 μm and 9 μm.

### Fluorescence *in situ* hybridization

Embryonic brains (E11.5 and E15.5) were fixed in 4% PFA in PBS at 4°C overnight and placed in 30% sucrose/PBS (weight/volume) until the tissue sank (12 h-16hrs). After cryopreservation, the brains were embedded in O.C.T. Compound (Sakura Finetek), frozen on dry ice, and stored at −80°C. Coronal brain sections (14 μm) were cut using a Leica CM3050S cryostat, mounted onto Superfrost Ultra Plus slides (Thermo Scientific), and stored at −20°C. Frozen sections were air-dried for approximately 30 min. For FISH analysis, RNAscope Fluorescent Multiplex Assays (ACD, 320850, 322000, and 322340) were conducted according to the manufacturer’s instructions (ACD, 320293-USM and 320535-TN) with RNAscope Probes directed against Fgf10 (ACD, 446371), Flrt1 (ACD, 555481) and Flrt3 (ACD, 490301). Immunostaining against CEP83 protein was performed prior to counterstaining with DAPI for Cep83 tKO embryos.

### Single-cell RNA sequencing and sample preparation

E15.5 of mutant embryos were removed from the uterus of mutant females, and stored in ice-cold in Leibowitz medium with 5% FBS. Brains were isolated and cut in 300 μm using a vibratome (Leica VT1000S, Germany) in cutting buffer, Leibowitz medium with 5% FBS. Then, brain sections were manually dissected for both hemisphere of cortical primordium of cortex. Meanwhile, genotyping was performed. Then, collected tissues were transferred into a dissociation Buffer, EBSS. Dissociation was performed with Worthington Kit and manually Papain dissociation system was carried out according to the recommended protocol (Worthington, #LK003163).

### Single-cell RNA library preparation

For experiments utilizing the 10x Genomics platform, the following reagents were used Chromium Single Cell Next GEM Single Cell 3’ GEM kit v3.1 (PN-1000130), Library Construction Kit (PN-1000196), Chromium Next GEM Single Cell 3′ Gel Bead Kit v3.1 (PN-1000129) and Single Index Kit T Set A (PN-1000213) and Dual Index Kit TT Set A (PN-1000215) were used according to the manufacturer’s instructions in the Chromium Single Cell 3′ Reagents Kits v3.1 User Guide.

### Sequencing

Transcriptome and barcode libraries were sequenced either on an Illumina NextSeq 500 and NovaSeq 6000 at the Next Generation Sequencing Facility of the Max Planck Institute of Biochemistry. Then, the sequencing data was processed with cellranger (version 7.0.1, reference ‘refdata-gex-mm10-2020’). Further processing was performed in R using Seurat (version 4.2.0) as follows: first, we merged the cellranger output files. We kept cells with more than 200 RNA molecules, less than 20% mitochondrial, more than 5% ribosomal, and less than 20% hemoglobin content. We used cellranger (version 7.0.1) to extract fastq files, align the reads to the mouse genome (10x genomics reference build MM10 2020 A), and obtain per-gene read counts. Subsequent data processing was performed in R using Seurat (version 4.2.0) with default parameters if not indicated otherwise. After merging the data, we normalized the data (normalization.method = ‘LogNormalize’, scale.factor = 10000), detected variable features (selection.method = ‘vst’, nfeatures = 2000), and scaled the data (vars.to.regress = c(’nCount_RNA’)). We then applied quality control filters on cells with the following criteria: a) more than 200 genes detected b) less than 20% mitochondrial genes reads c) more than 5% ribosomal protein genes reads d) less than 20% hemoglobin genes reads e) singlets as determined by doubletFinder (version 2.0.3, pK = 0.09, PCs=1:10). After performing principal component analysis on variable features, nearest-neighbor graph construction and UMAP dimension reduction were carried out on PCs 1-20, followed by cell clustering at a resolution of 0.2. Neuronal subset clustering was performed with are resolution of 0.5.

### Computational simulations

We modelled the attraction forces for FLRT-positive, -mutant and -negative cells as sine curves based on *in vivo* gain- and loss-of-expression experiments as previously described (Seiradake 2014, del toro 2017). The equation had the following terms: Attraction curve = κ[Asin(λx)]+bs+ε, where A is the amplitude and λ the frequency. The frequency value was the same for both attraction curves (λ, 0.051) but not their amplitude (A, FLRT-positive and mutant: 41,41, FLRT-negative:56.74). The strength of both curves was adjusted with the term κ to represent both: the balanced situation with slightly higher attraction for FLRT-positive particles (k, FLRT-positive: 1/4, FLRT-negative: 1/10), and the Flrt1/3 dKO situation with low attraction between FLRT-mutant particles and strong attraction between FLRT-negative particles (k, FLRT-positive: 1/10, FLRT-negative: 1/2).The basal subtraction value (bs, 100) was used for fitting both curves, factor ε changes randomly from −10 to 10 and is used to add noise.

We used particles to represent FLRT-positive, -mutant and FLRT-negative cells using the particle system toolbox, MATLAB particles version 2.1^52^, as previously described (del Toro 2017). FLRT-positive or -mutant particles were arranged in an 8 by 36 matrix, spaced by 0.2 units, and were given an attraction towards neighboring particles in both axes using their attraction curve. FLRT-negative particles were arranged in a matrix shifted 0.1 units on both the X and Y axes relative to the previous matrix (50:50 condition) and were attracted to neighboring particles in both axes using their attraction curve.The number of particles in both matrices was adjusted independently to mimic the overall increase in particles and the FLRT-mutant to FLRT-negative ratio observed in experiments with tKO mice.

All particles received random speeds (ranging from 6 to 12 arbitrary units) for moving along the Z axis and were simulated for 100 frames (0.001 units step time). After simulation, the position of every particle was retrieved, and their area was calculated by triangulating the surface formed by their positions and then summing the areas of the triangles. The 3D surface and 2D contour lines representing the central slice of the surface plot at the median Y value were plotted in MATLAB.

### Quantification and Statistical analysis

Images were processed with the open-source image analysis software Fiji. Automatic cell counting analysis was performed using open-source CellProfiler 4.2.5 and software Fiji with plugin ‘Cell counter’. Kernel Density Estimation (Heat map) was performed using Rstudio, Statistical significance was determined using one-way ANOVA Tukey’s post hoc test or t test with Welch correction the using Prism version 9,10 (Graphpad Software). Statistical significance was defined as p < 0.05. All values in the text and in the figure legends indicate mean +/− SEM.

## Extended data figure legends

**Figure S1.**
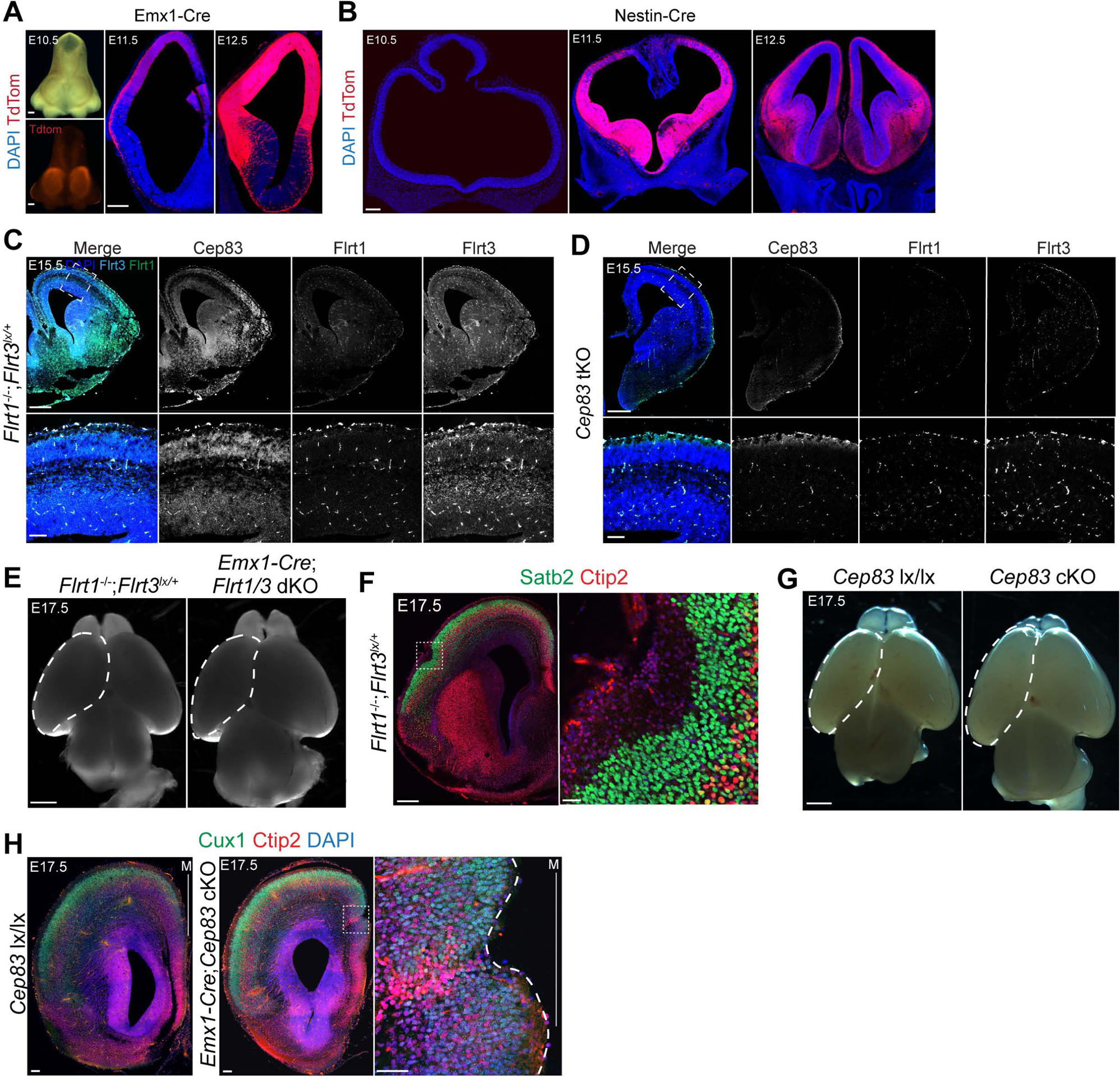
related to Figure 1 Loss of Cep83 and Flrt1/Flrt3 enhances cortical folding and the presence of sulci. (A) Macroscopic image of E10.5 of Emx1-Cre embryo, head only (above). tdTomato expression of Emx1-Cre in embryo brain (below). The Cre-dependent reporter pCALNL-TdTom expression with DAPI in E11.5 and E12.5 of coronal view of cortex. (B) The Cre-dependent reporter pCALNL-TdTom expression tdTomato expression of Nestin-Cre with DAPI in E10.5, E11.5 and E12.5 of coronal view of cortex. (C) Double *in situ* hybridization (ISH) for Flrt1 and Flrt3 combined with Cep83 antibody staining in coronal sections of E15.5 cortex of *Flrt1*^−/−^;*Flrt3*^lx/+^. Area in dashed rectangle is shown with higher magnification on the right. (D) Double ISH for Flrt1 and Flrt3 combined with Cep83 antibody staining in coronal sections of E15.5 cortex of *Cep83* tKO. Area in dashed rectangle is shown with higher magnification on the right. (E) Representative whole-mount images of E17.5 *Flrt1* ^−/−^*Flrt3* ^lx/+^ and Emx1-Cre;*Flrt1/3* dKO brains. Dashed areas were measured to obtain quantification in Figure1 A. (F) E17.5 *Flrt1*^−/−^;*Flrt3*^lx/+^ brain section labeled with Satb2 (green), Ctip2 (red). A folding area in dashed rectangle is shown with higher magnification on the right. (G) Representative whole-mount images of E17.5 *Cep83* lx/lx and *Cep83* cKO brains. Dashed areas were measured to obtain quantification in Figure 1 C. (H) E17.5 *Cep83* lx/lx and Emx1-Cre;*Cep83* cKO brain sections labeled by Cux1 (green), Ctip2 (red), and DAPI (blue). Area in dashed rectangle is shown with higher magnification on the right. The midline is indicated with a vertical line and the letter ‘M’. Scale bars represent 20 μm, 20 μm, 100 μm (A), 100 μm (B), 500 μm, 100 μm. (C), 500 μm, 100 μm. (D), 1mm (E), 1mm, 100 μm (F), and 1 mm (G), 100 μm, 100 μm, 100 μm (H).

**Figure S2.**
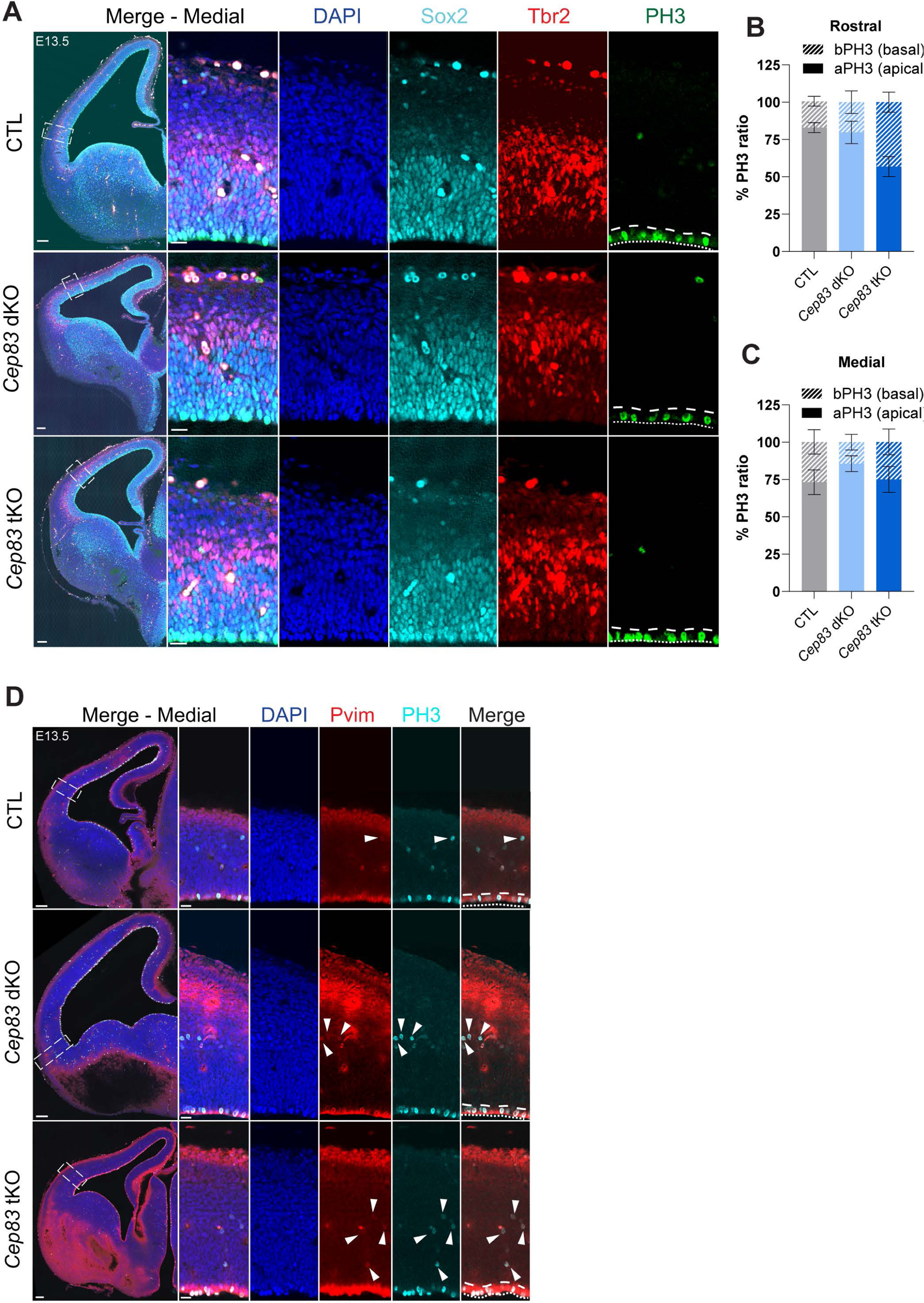
related to Figure 2 *Cep83* tKO mice have more intermediate progenitors. (A) E13.5 medial cortices of CTL (*Flrt1*^−/−^;*Flrt3*^lx/+^), *Cep83* dKO. and *Cep83* tKO embryos were stained with DAPI (blue), Sox2 for early neuronal progenitors (cyan), Tbr2 for intermediate progenitors (red), and PH3 for mitotic cells (green). The apical side of the VZ is indicated with a dotted line, the basal side with a dashed line in PH3 stained images. Areas in dashed rectangles in (A, left row) are shown with higher magnification on the right. (B) Proportion of apical/basal mitotic cells (PH3) in rostral region (CTL (*Flrt1*^−/−^;*Flrt3*^lx/+^) n = 5 brains, *Cep83* dKO n = 5 brains, *Cep83* tKO, n = 5 brains). Data are shown as mean ± SEM; aPH: CTL vs *Cep83* dKO, p = 0.486, CTL vs *Cep83* tKO, p = 0.093, *Cep83* dKO vs *Cep83* tKO, p = 0.516, bPH: CTL vs *Cep83* dKO, p = 0.515, CTL vs *Cep83* tKO, p = 0.101, *Cep83* dKO vs *Cep83* tKO, p = 0.515. one-way ANOVA with Tukey’s post hoc analysis. (C) Proportion of apical/basal mitotic cells (PH3) in medial region (CTL (*Flrt1*^−/−^;*Flrt3*^lx/+^) n = 5 brains, *Cep83* dKO n = 5 brains, *Cep83* tKO, n = 5 brains). Data are shown as mean ± SEM; aPH: CTL vs *Cep83* dKO, p = 0.497, CTL vs *Cep83* tKO, p = 0.985, *Cep83* dKO vs *Cep83* tKO, p = 0.595, bPH: CTL vs *Cep83* dKO, p = 0.481, CTL vs *Cep83* tKO, p = 0.984, *Cep83* dKO vs *Cep83* tKO, p = 0.579. one-way ANOVA with Tukey’s post hoc analysis. (D) E13.5 medial cortices of CTL (*Flrt1*^−/−^;*Flrt3*^lx/+^), *Cep83* dKO. and *Cep83* tKO embryos were stained with DAPI (blue), Pvim for dividing RG cells (red), PH3 for mitotic cells (cyan). The apical side of the VZ is indicated with a dotted line, the basal side with a dashed line in Pvim/PH3 images. The basal progenitors are present as Pvim/PH3 co-immunopositive cells located above apical side of the VZ (marked by arrow heads). Scale bars represent 100 μm, 50 μm (A), and 100 μm, 20 μm (D)

**Figure S3.**
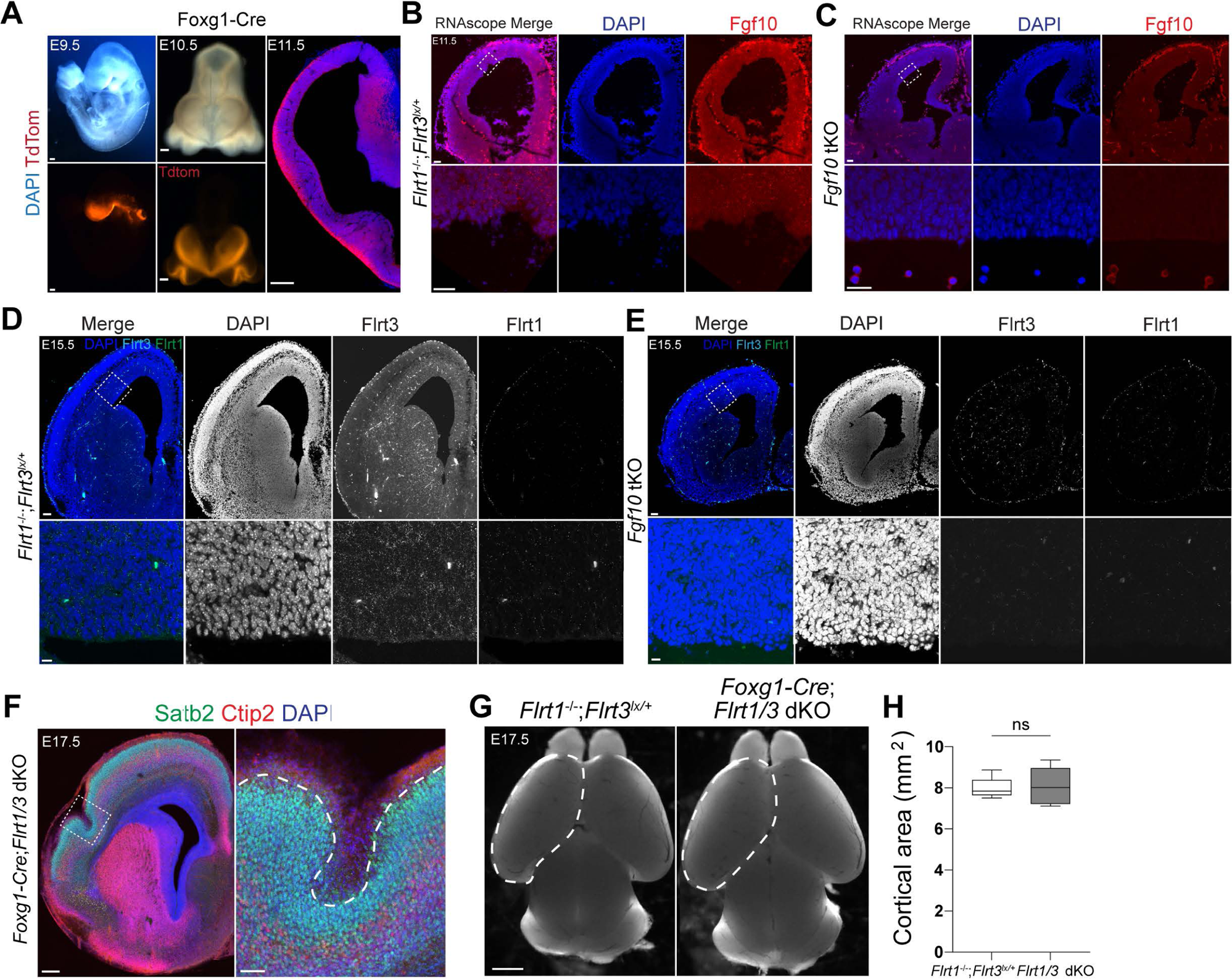
related to Figure 3 Loss of Fgf10 and Flrt1/Flrt3 enhances cortical folding and the presence of gyri. (A) Macroscopic image of E9.5, E10.5 and E11.5 of Foxg1-Cre embryo. E9.5 whole embryo and E10.5 head only (above) is shown with the Cre-dependent reporter pCALNL-TdTom expression of Foxg1-Cre in embryo brain (below) and with DAPI in E11.5 of coronal view of cortex. (B) ISH for Fgf10 in coronal sections of E11.5 cortex of *Flrt1*^−/−^;*Flrt3*^lx/+^. Area in dashed rectangle is shown in higher magnification of the VZ. (C) ISH for Fgf10 in coronal sections of E11.5 cortex of *Fgf10* tKO. Area in dashed rectangle is shown in higher magnification of the VZ. _(D)_ Double ISH for Flrt1 and Flrt3 with DAPI in coronal sections of E15.5 cortex of *Flrt1*^−/−^;*Flrt3*^lx/+^. Area in dashed rectangle is shown with higher magnification on the right. (E) Double ISH for Flrt1 and Flrt3 with DAPI in coronal sections of E15.5 cortex of *Fgf10* tKO. Scale bars, Area in dashed rectangle is shown with higher magnification on the right. (F) Representative E17.5 *Foxg1-Cre*;*Flrt1/3* dKO brain section labeled with Satb2 (green), Ctip2 (red), and DAPI (blue). Area in dashed rectangle is shown with higher magnification on the right. Dashed line indicate a gyrus. (G) Representative whole-mount images of E17.5 *Flrt1* ^−/−^*Flrt3* ^lx/+^ and *Foxg1-Cre*;*Flrt1/3* dKO brains. Dashed area were measured to obtain quantifications. (H) Quantifications of the cortical areas shown in (G). *Flrt1* ^−/−^*Flrt3* ^lx/+^, n = 10 brains; Foxg1-Cre;*Flrt1/3* dKO, n = 11 brains. p = 0.859, unpaired t-test with Welch correction. Scale bars represent 100 μm, 20 μm, 100 μm, 20 μm, 100 μm (A), 50 μm, 25 μm (B), 50 μm, 25 μm (C), 100 μm, 10 μm (D), 100 μm, 10 μm (E), 200 μm, 50 μm (F), and 1 mm (G).

**Figure S4.**
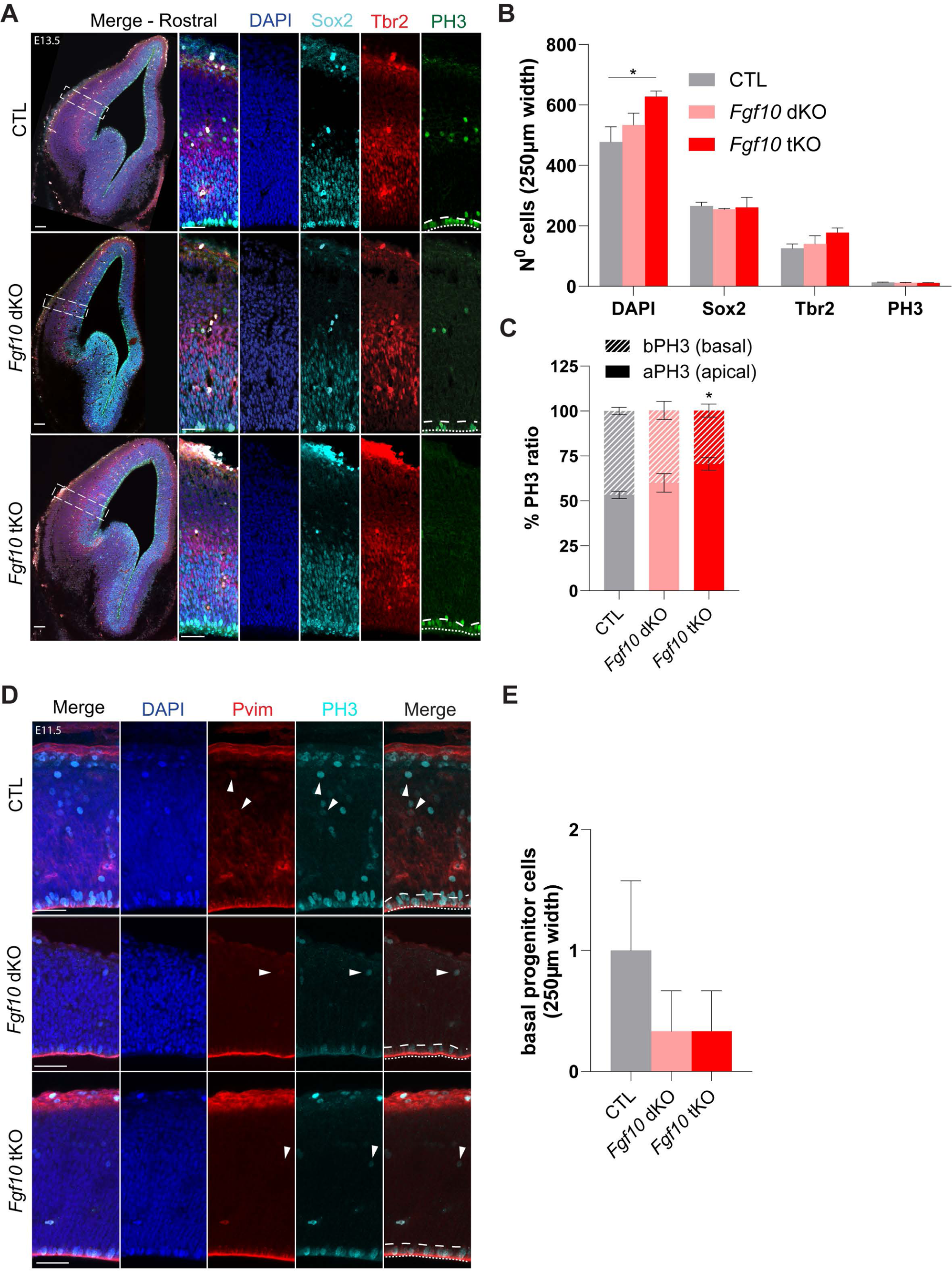
related to Figure 4 Progenitor expansion in *Fgf10* tKO *and Fgf10* dKO embryos at E11.5. (A) E13.5 rostral cortices of CTL (*Flrt1*^−/−^*;Flrt3* ^lx/+^), *Fgf10* dKO and *Fgf10* tKO embryos were stained DAPI (blue), apical progenitors Sox2 (cyan), intermediate progenitors Tbr2 (red), and mitotic cells PH3 (green). Apical and basal sides of the VZ are indicated with dotty and dashed lines in PH3 stained images, respectively. Areas in dashed rectangles in (A) are shown with higher magnification on the right. (B) Quantification of the data shown in (A). CTL (*Flrt1*^−/−^;*Flrt3*^lx/+^), n = 4 brains, *Fgf10* dKO n = 3 brains, *Fgf10* tKO n = 4 brains. Data are shown as mean ± SEM; Data are shown as mean ± SEM; CTL vs *Fgf10* tKO p = 0.048, *p < 0.05, one-way ANOVA with Tukey’s post hoc analysis. (C) Proportion of apical/basal mitotic cells (PH3) in rostral region CTL (*Flrt1*^−/−^;*Flrt3*^lx/+^), n = 3, *Fgf10* dKO n = 3, *Fgf10* tKO n = 4. Data are shown as mean ± SEM; aPH: CTL vs *Fgf10* dKO, p = 0.827, CTL vs *Fgf10* tKO p = 0.076, *Fgf10* dKO vs *Fgf10* tKO p = 0.182. bPH: CTL vs *Fgf10* dKO, p = 0.537, CTL vs *Fgf10* tKO p = 0.037, *Fgf10* dKO vs *Fgf10* tKO p = 0.185. *p < 0.05, one-way ANOVA with Tukey’s post hoc analysis. (D) E11.5 rostral cortices of CTL (*Flrt1*^−/−^;*Flrt3*^lx/+^), *Fgf10* dKO. and *Fgf10* tKO embryos were stained with DAPI (blue), Pvim for dividing RG cells (red), PH3 for mitotic cells (cyan). The apical side of the VZ is indicated with a dotted line, the basal side with a dashed line in Pvim/PH3 images. The basal progenitors are present as Pvim/PH3 co-immunopositive cells located above apical side of the VZ (marked by arrow heads). _(E)_ Quantifications of cell densities in the rostral regions shown in (D).. CTL (*Flrt1*^−/−^;*Flrt3*^lx/+^), n = 3 brains,*Fgf10* dKO n = 3 brains, *Fgf10* tKO, n = 3 brains). Data are shown as mean ± SEM; p = 0.551 (CTL vs Fgf10 dKO/tKO), no significant changes between groups, one-way ANOVA with Tukey’s post hoc analysis. Scale bars represent 100 μm, 50 μm (A), and 50 μm (D)

**Figure S5.**
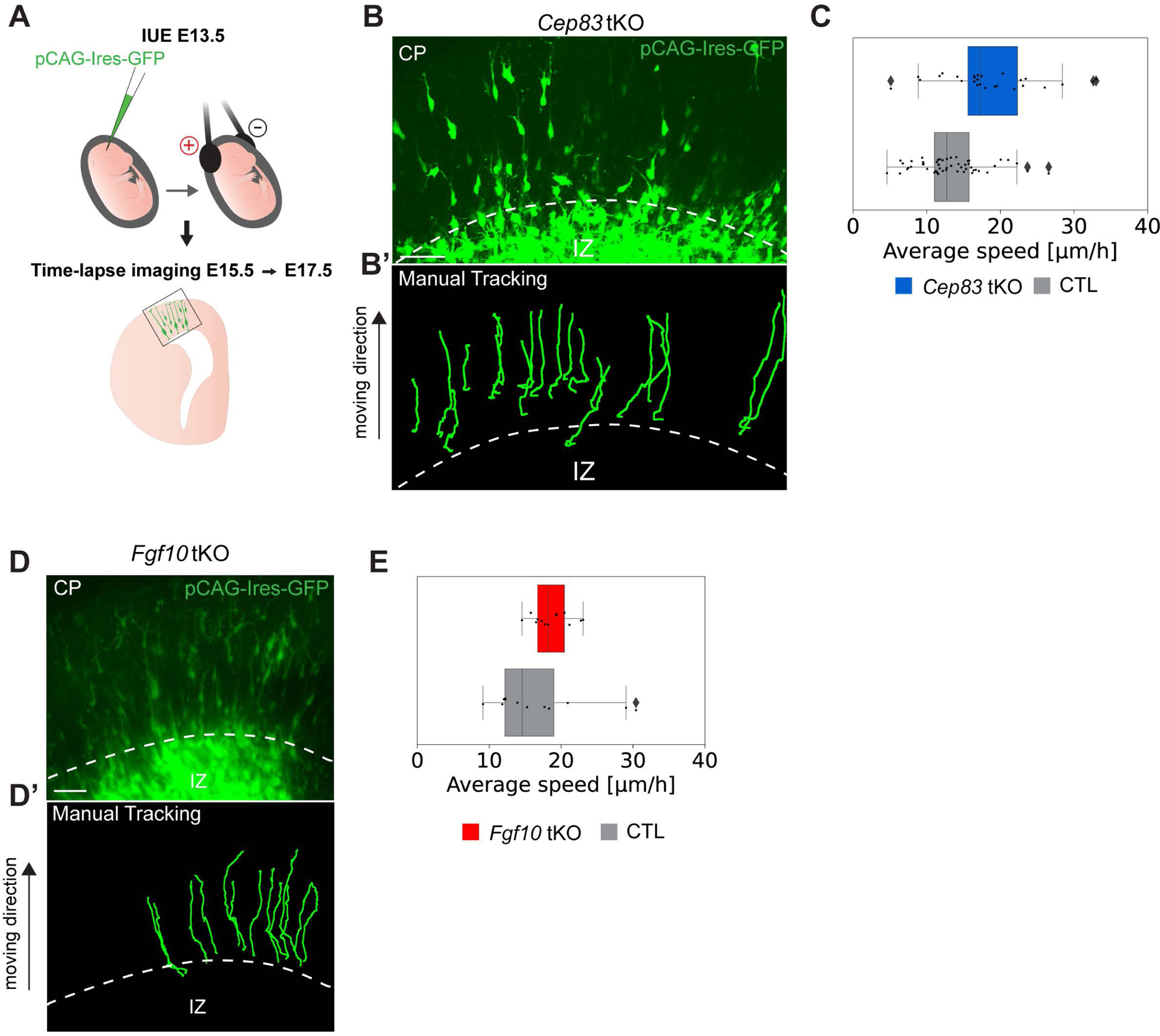
related to Figure 5 Higher cell density in gyri of *Fgf10* tKO mice. (A) Schematic diagram of *in utero* electroporation (IUE) performed at E13.5 of embryo.. The time-lapse imaging starts at E15.5 with ex vivo sliced section labeled by pCAG-Ires-GFP by IUE over 48 hours. (B) Migrating GFP labelled neurons by IUE were tracked in the CP in ex vivo sliced section of *Cep83* tKO. (B’) Migration paths of manually tracked neurons within CP for speed analysis. _(C)_ Quantification of the tracked neurons in (B’). The average speed of tracked neurons is represented as a box plot, with median (centre line), 50% interquartile range (box) and whiskers extending 1.5 times the interquartile range; Average speed values of individual neurons are represented as a scatterplot, *Cep83* tKO, n = 2, CTL (*Flrt1*^−/−^;*Flrt3*^lx/+^), n = 3. (D) Migrating GFP labelled neurons by IUE was tracked in the CP in ex vivo sliced section of *Fgf10* tKO. (D’) Progression lines of tracked migrating neurons within CP for speed analysis. (E) Quantification of tracked neurons in (D’). The average speed of tracked neurons is represented as a box plot, with median (centre line), 50% interquartile range (box) and whiskers extending 1.5 times the interquartile range; Average speed values of individual neurons are represented as a scatterplot; *Fgf10* tKO, n =1, CTL (*Flrt1*^−/−^;*Flrt3*^lx/+^), n = 1. Scale bars represent 50 μm (B), and 50 μm (D).

**Figure S6.**
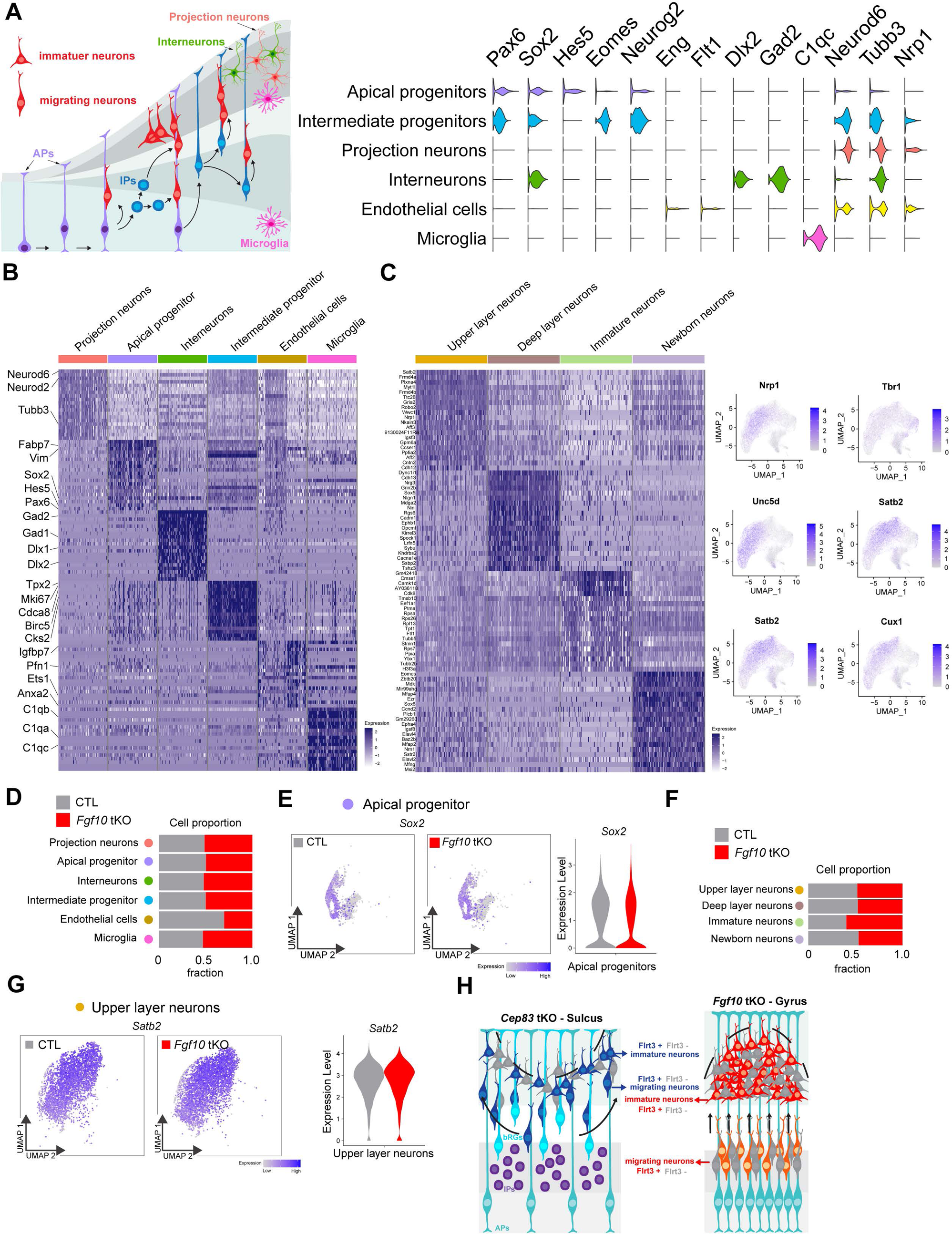
related to Figure 6 Modeling cortex folding in *Cep83* tKO and *Fgf10* tKO mice. (A) Expression of marker genes per cell type. Cell types are grouped on the basis of their identity and shared maker genes. (B) Gene signatures for all cell types identified in the combined all genotypes. Top 30 differentially expressed genes for each cell type are presented. Cells were down-sampled to a maximum of 100 cells per cell type (C) Gene signatures for all cell types identified in the combined all genotypes. Top 30 differentially expressed genes for each cell type are presented. Cells were down-sampled to a maximum of 100 cells per cell type Expression of canonical marker genes for selected cell types in the UMAP visualization of the combined all genotypes. (D) Proportion of each cell type by genotype in CTL (*Flrt1*^−/−^;*Flrt3*^lx/+^) and *Fgf10* tKO mice. (E) Expression of the apical progenitor marker gene Sox2 in apical progenitor cell type for CTL (*Flrt1*^−/−^*;Flrt3*^lx/+^) and *Fgf10* tKO embryos. (UMAP and violin plot). (F) Proportion of sub cell type by genotype in CTL (*Flrt1*^−/−^;*Flrt3*^lx/+^) and *Fgf10* tKO mice. _(G)_ Expression of CP layer marker gene, Satb2, in upper layer neurons for CTL (*Flrt1*^−/−^*;Flrt3*^lx/+^) and *Fgf10* tKO embryos. (UMAP and violin plots). (H) Graphic summary of how folds are developed in *Cep83* tKO (Sulcus) and *Fgf10* tKO(Gyrus)

